# Coincidence detection supported by electrical synapses is shaped by the D-type K^+^ current

**DOI:** 10.1101/2025.09.17.676857

**Authors:** Antonella Dapino, Sebastián Curti

**Affiliations:** Laboratorio de Neurofisiología Celular, Unidad Académica de Fisiología, Facultad de Medicina, Universidad de la República, Montevideo 11800, Uruguay

## Abstract

Electrical synapses mediated by gap junctions are widespread in the mammalian brain, playing essential roles in neural circuit function. Beyond their role synchronizing neuronal activity, they also support complex computations such as coincidence detection—a circuit mechanism in which differences in input timing are encoded by the firing rates of coupled neurons, enabling preferential responses to synchronous over temporally dispersed inputs. Electrical coupling allows each neuron to act as a current sink for its partner during independent depolarizations, thereby reducing excitability. In contrast, synchronous inputs across the network minimize voltage differences through gap junctions, reducing current shunting and increasing spiking probability. However, the contribution of intrinsic neuronal properties to coincidence detection remains poorly understood. Here, we investigated this issue in the Mesencephalic Trigeminal (MesV) Nucleus of mice, a structure composed of somatically-coupled neurons. Using whole-cell recordings and pharmacological tools, we examined the role of the D-type K+ current (I_D_), finding that it critically shapes both the intrinsic electrophysiological properties of MesV neurons and the dynamics of electrical synaptic transmission. Its fast activation kinetics and subthreshold voltage range of activation make I_D_ a key determinant of transmission strength and timing. Furthermore, the I_D_, likely mediated by Kv1 subunits, is expressed at the soma and the axon initial segment. Finally, we characterized two key parameters of coincidence detection—precision (time window for effective input summation) and gain (differential response to coincident versus dispersed inputs)—finding that I_D_ enhances precision by accelerating membrane repolarization and reduces the gain by limiting neuronal excitability.

**SUMMARY:** Electrical synapses enable neural circuits to perform as coincidence detectors, that is, to preferentially respond to simultaneous inputs. The temporal precision of this operation is critically determined by voltage-dependent conductances of the soma and proximal axon.

## INTRODUCTION

Electrical synaptic transmission is a modality of communication based on the direct flow of current between neurons typically mediated by gap junctions, that supports the synchronic activation of neuronal ensembles given its fast and bidirectional nature (Bennett and Zukin, 2004; Connors and Long, 2004; Perez Velazquez and Carlen, 2000). Nonetheless, electrical synapses can strongly reduce the excitability of all neurons of the circuit. During synaptic inputs to a given neuron, part of the undelaying current flows into the electrically coupled partners. As a result, gap junctions act as current sinks, reducing the input resistance of all neurons within the circuit. This effect, known as “loading”, reduces the efficacy of depolarizing inputs and their ability to induce action potential firing (Getting and Willows, 1974; Bennett and Zukin, 2004; Alcami and Marty, 2013; Alcami, 2018; Davoine and Curti, 2019). In contrast, synchronous inputs to all electrically coupled neurons reduce the voltage drop across junctions, minimizing the current leak; a condition termed “unloading”. In this way, by reducing changes in membrane potential to temporally distributed depolarizations, electrical synapses endow neural circuits with the ability to selectively respond to synchronic inputs, thereby promoting coincidence detection (Curti et al., 2012; Galarreta and Hestrin, 2001; Veruki and Hartveit, 2002; Rela and Szczupak, 2003; Curti et al., 2022; Trigo et al., 2025).

Mesencephalic trigeminal (MesV) neurons are unique among primary afferents in that their cell bodies are located within the brainstem (Weinberg, 1928). They innervate muscle spindles of jaw-closing muscles and mechanoreceptors in the periodontal ligaments (Gottlieb et al., 1984). Centrally, their axons project to key motor and premotor centers, including the trigeminal motor nucleus, the rostral parvocellular reticular formation, and the nucleus supratrigeminalis (Dessem and Taylor, 1989; Liem et al., 1991). In addition to their sensory role, MesV neurons receive input from the hypothalamus and various brainstem nuclei (Lazarov, 2002; Verdier et al., 2004), supporting the notion that they serve dual functions as primary afferents and interneurons within central circuits involved in the generation of orofacial behaviors (Morquette et al., 2012). MesV neurons are electrically coupled in pairs or small clusters via large somatic gap junctions containing connexin36 (Curti et al., 2012). Electrical transmission between these afferents is modulated by the neurons’ intrinsic electrophysiological properties, which play a critical role in regulating coincidence detection (Davoine and Curti, 2019). In previous work, we identified the D-type potassium current (I_D_) as a key determinant of species-specific differences in the function of electrical synapses between rats and mice (Dapino et al., 2023). Notably, although coupling strength is comparable, electrical transmission in rats promotes lateral excitation and synchronization between MesV neuron pairs, phenomena that are rarely observed in mice. This disparity is attributed to a markedly higher expression of the I_D_ in mice, which not only reduces neuronal excitability but also the amplitude and duration of postsynaptic potentials, given its fast activation kinetics and its subthreshold voltage range of activation (Dapino et al., 2023). These findings highlight the critical role of I_D_ in shaping the electrophysiological properties of MesV neurons and regulating electrical synapse function.

Building on this foundation, the present study investigates the specific contribution of I_D_ to electrical transmission and coincidence detection in mice MesV neurons. We demonstrate that I_D_ affects both passive and active membrane properties. Its steady-state activation decreases input resistance (Rin) and attenuates coupling strength. During spike-evoked coupling, rapid I_D_ activation diminishes the amplitude and duration of postsynaptic potentials while accelerating their time to peak, thereby enhancing precision of transmission. Furthermore, we show that I_D_ is expressed in both the soma and proximal axon, which are strongly electrotonically coupled, suggesting dynamic bidirectional interactions between these compartments. Finally, we provide evidence that I_D_ regulates both the temporal window and gain of coincidence detection, enhancing precision at the cost of reduced gain.

## MATERIALS AND METHODS

### Ethics approval

Mice C57/BL (age P12-P17), of either sex, were obtained from the university animal facility (Unidad de Reactivos para Biomodelos de Experimentación, Facultad de Medicina), accredited by the university authorities (CHEA, Comisión Honoraria de Experimentation Animal). All animal care and experimental procedures were performed under the national guidelines and laws, with minimization of the number of animals used. Experimental protocol (N° 070151-000052-22) was approved by the School of Medicine ethics committee (CEUA, Comisión de Ética en el Uso de Animales) and national animal regulation authorities (CNEA, Comisión Nacional de Experimentación Animal).

### Slice preparation and electrophysiological recordings

Mice were decapitated without anesthesia, and their brains were quickly removed. Transverse brainstem slices (200 μm) were obtained by vibratome (DSK DTK-1000) while submerged in cold sucrose solution (248 mM sucrose, 26 mM HCO_3_, 10 mM glucose, 2.69 mM KCl, 2 mM MgSO_4_,1.25 NaH_2_PO_4_, 1mM CaCl_2_, 0.35 mM ascorbic acid, 0.3 mM pyruvic acid). Afterwards, slices were transferred into an incubation chamber filled with artificial cerebrospinal fluid solution (aCSF; 124 mM NaCl, 26 mM HCO_3_, 10 mM glucose, 2.69 mM KCl, 2 mM MgSO_4_, 2 mM CaCl_2_, 1.25 mM KHPO_4_) at 34°C for 30 min. Then, slices were kept at room temperature in aCSF until transferred into the recording chamber. Both sucrose and aCSF solution were bubbled permanently with carbogen (O_2_ 95% and 5% CO_2_) to maintain pH ∼7.4.

The recording chamber was mounted on an upright microscope stage (E600; Nikon EclipseFNI) and continuously perfused with aCSF (2 ml/min) at constant room temperature (25°C).

Whole-cell patch recordings were performed under visual control using infrared differential interference contrast (IR-DIC) equipped with a CMOS camera (ThorLabs).

MesV neurons are identified due to their location, morphological and electrophysiological characteristics (Curti et al., 2012). Recording pipettes pulled from borosilicate glass (AM-systems and WPI, 4-10 MΩ) were filled with intracellular solution (148 mM K-gluconate, 10 mM HEPES 4 mM, Na_2_-ATP, 3 mM MgCl_2_, 0.3 mM Na-GTP, and 0.2 mM EGTA, with pH ∼7.2). For studying electrical synaptic transmission, whole-cell recordings at soma were done simultaneously in two neurons. For studying soma-axon coupling, a neuron was patched at the soma (with pipette filled with Alexa Fluor^TM^ 488, to visualize the axon cut end, “bleb”) and with a smaller pipette (resistance ∼10-15 MΩ) at the axon, to obtain simultaneous recordings.

Seal resistance between the microelectrode tip and the cell membrane was > 1 GΩ, and membrane capacitance was compensated before breaking the seal by negative pressure suction. Recordings were made by using Multiclamp 700B amplifier (Molecular Devices), low-pass filtered at 5kHz, sampled at 10-40 kHz (depending on the experiment), and acquired by means of an analog-to-digital convertor (National Instruments, NI USB- 6221) connected to a computer. In current clamp configuration, the voltage-drop across the microelectrode resistance was eliminated by bridge balance control of the amplifier; while in voltage clamp configuration, both membrane capacitance and series resistance were continuously monitored and compensated (80%). Healthy cells included in this study were identified by fulfilling 2 out of the 3 following criteria: input resistance (Rin) > 60 MΩ, resting membrane potential (RMP) < -50 mV, and action potential amplitude > 60 mV.

### Pharmacology

Drugs were applied either by addition to the bath solution or by localized pressure ejection (puff) for experiments aimed to assess I_D_ localization. Localized applications were performed using a Picospritzer (5–10 psi) connected to a patch pipette with a tip diameter of ∼2 μm. 4-aminopyridine (4-AP; Sigma) and α-dendrotoxin (α-DTX; Alomone) were prepared in aCSF at their final concentrations; α-DTX solutions additionally contained 0.1% bovine serum albumin (BSA; Sigma).

### I_D_ recording

To reduce current intensity through voltage-gated Na+ channels and HCN channels, NaCl of the aCSF was substituted by choline-Cl (Sigma), and TTX (0.25 μM, Sigma) and CsCl (2.5 mM, Sigma) were added. In this condition (control), a series of step-like voltage commands of 500 ms in duration were applied, from 0 to 70 mV in steps of 5 mV starting from a holding potential of −70 mV. These voltage commands were repeated after the addition of different concentrations of 4-AP. For dose-response curves (**Fig. S1**), increasing concentrations of 4-AP were added into the aCSF (0.03, 0.3, 3, 30, 60, 100, 300 μM), and the I_D_ was isolated by subtracting current traces obtained after the addition of 4-AP from those obtained in control. Activation curves for obtaining maximum conductance values and half-activation voltage for each concentration were constructed as described previously (Dapino et al., 2023).

### Epifluorescence

Experiments where MesV neuron’s axon tracing were required, Alexa Fluor^TM^ 488 NHS ester (50 μM, ThermoFisher) was added to the intracellular solution. 5-10 min after whole-cell configuration was established, the axon trajectory was examined under the fluorescent microscope mentioned previously, equipped with a X40 water-immersion objective (NA 0.8) and LED light (pE-300, CoolLED) at power 5-20%. Images were acquired with ThorCam software. To position the puff pipette near the axon, switching back and forth between the epifluorescent and DIC images was needed, minimizing the fluorescence exposure time to avoid cell damage. No changes in the electrophysiological properties of MesV neurons were detected under these conditions.

### Input resistance (Rin) measurement

In current clamp, a series of hyperpolarizing current pulses from -450 to 0 pA were applied in steps of 50 pA, and the voltage responses (Vm) were recorded. Plots of Vm change (at the peak of the hyperpolarizing response, previous to the sag which indicates HCN current activation) as a function of the intensity of the injected current were constructed and data was fitted by a simple linear regression. Rin was determined as the slope of the regression.

### Coupling coefficient (CC) measurement

During simultaneous current clamp recordings of pairs of MesV neurons, a series of hyperpolarizing current pulses from -450 to 0 pA in steps of 50 pA were applied to one neuron (presynaptic), and the voltage response in the injected neuron (Vm pre) and the coupled one (postsynaptic, Vm post) were recorded. Plots of Vm post as a function of Vm pre were constructed and data was fitted by a linear function, whose slope represents the CC. Vm pre and Vm post are measured at the peak of corresponding hyperpolarizing responses. CC is reported as directions, two directions per coupled pair. In the case of CC_spike_, to study spike transmission efficacy, it was defined as the ratio between the peak amplitude of the postsynaptic spikelet and that of the presynaptic spike which elicited the spikelet.

### Gap junction conductance (Gj) measurement

During simultaneous voltage clamp recordings from pairs of MesV neurons, voltage commands of increasing amplitude in steps of -5 mV were applied into one of the neurons from a holding potential of -55 mV up to -105 mV, while the other neuron was held at -55 mV. The membrane current in the non-stepped cell, corresponding to the junctional (synaptic) current (Ij), was measured. From these recordings, plots of the Ij as a function on the transjunctional voltage (membrane voltage difference between cells during commands) were constructed, and the Gj was estimated from the slope of linear regressions. Gj is reported as directions, two directions per coupled pair.

### Assessment of MesV neurons excitability (firing gain)

During current clamp recordings, a series of depolarizing current pulses of 200 ms in duration were applied, whose intensities ranged from 0 to 900 pA, in steps of 50 pA. From these recordings, plots of the spike count versus current intensity were constructed and the firing gain, defined as the slope of linear regression forced through the origin, was determined. This parameter encompasses the ability of the neuron to produce repetitive discharges as well as its rheobase (minimal current for eliciting firing), representing a valuable indicator of neuronal excitability (Dapino et al., 2023).

### Estimation of firing level (threshold)

From action potential recordings in current clamp, the firing level was defined as the membrane voltage value at which the first derivative of the action potential first reaches values > 10 mV/ms in a monotonic increasing range.

### Estimation of membrane time constant

A brief current pulse (-50 pA, 10 ms; in order to minimize activation or deactivation of any voltage-gated current) was applied at RMP, and the voltage response was recorded. To estimate the membrane time constant (𝜏m), the onset of the membrane voltage response was fitted to an exponential function of the form:

### Image processing

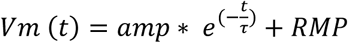

IR-DIC and epifluorescence images were improved using ImageJ by enhancing contrast (0.5-1% saturated pixels) and brightness/contrast balancing. For measuring axonal length, Z-projection of epifluorescence images stacks were created, and axonal length was measured from axon hillock up to the axonal bleb or until fluorescence was lost due to axonal penetration into the brain slice.

### Data analysis

Data were analyzed by using IgorPro9 (WaveMetrics), Graphpad 8, and Python scientific development environment Spyder (libraries: Axographio, Lmfit, Matplotlib, Numpy, Pandas, Scipy, and Stfio). Results were expressed as average value ± standard deviation [SD]. The significance of quantitative data was determined by using Graphpad Prism 8 and IgorPro9 statistical analysis. Statistical significance in figures is denoted using asterisk notation as follows: p <0.05 (*), p <0.01 (**), p <0.001 (***), and p <0.0001 (****).

## RESULTS

### Contribution of I_D_ to intrinsic properties of MesV neurons

MesV neurons could exhibit strong electrical coupling; however, action potential firing rarely triggers activation of its coupled partners in mice. This is attributed to the intrinsically low excitability of these neurons, resulting from their high expression of the I_D_ (Dapino et al., 2023). These observations support the idea that the I_D_ plays a crucial role in shaping the electrophysiological properties of MesV neurons and, consequently, in modulating the functional impact of electrical synaptic transmission. To explore this further, we first examined the contribution of the I_D_ to the intrinsic electrophysiological properties of these afferents. To this aim we employed 4-aminopyridene (4-AP), a broad-spectrum K^+^ voltage-gated channel blocker that, at micromolar concentrations, selectively targets the I_D_. However, complete blockade of this conductance typically induces spontaneous firing and destabilizes the membrane potential, thereby preventing reliable measurement of membrane properties. To address this limitation, we empirically determined the highest concentration of 4-AP that did not induce spontaneous activity (10 μM). Dose-response curves rendered an IC50 that averaged 4.45 ± 2.69 μM [SD], whereas 10 μM results in a blockade of about 75% of the total I_D_ (**Fig. S1 A-F**). Noteworthy, activation curves of membrane currents sensitive to the tested concentration range (0.03 to 300 μM) showed that the half-activation voltage (Vhalf) did not present any statistical difference, confirming that within this concentration range 4-AP selectivity target the I_D_ (**Fig. S1 G**; Vhalfs [0.3 μM]: -26.05 ± 9.04, n = 9 cells, N = 5 mice; [3 μM]: -32.24 ± 5.72, n = 9 cells, N = 5 mice; [30 μM]: -33.24 ± 3.18, n = 13 cells, N = 6 mice; [60 μM]: -32.06 ± 2.63, n = 6 cells, N = 3 mice; [100 μM]: -31.72 ± 2.81, n = 7 cells, N = 3 mice; [300 μM]: -30.33 ± 2.95, n = 7 cell, N = 3 mice; p = 0.0512, one-way ANOVA).

Bath applications of 10 μM 4-AP resulted in an enhancement of excitability of MesV neurons (**Fig. 1 A**). This was evidenced by a significant increase in firing gain, defined as the slope of the spike-current relationships (see Materials and Methods), which increased from 3.19 ± 4.67 spikes/nA [SD] in control, to 10.50 ± 15.02 spikes/nA [SD] following 4-AP application (n = 57 cells, N = 17 mice; p = 2.27 × 10⁻⁵, paired two-tailed t-test; **Fig. 1 B-C**). This indicates that the I_D_ critically contributes to the active membrane properties of MesV neurons by markedly suppressing neuronal excitability. These changes were accompanied by depolarization of the resting membrane potential (RMP; **Fig. 1 D**) (Control: -53.12 ± 3.72 mV [SD]; 4-AP: -50.01 ± 3.13 mV [SD]; n = 45 cells, N = 11 mice; p = 3.85 × 10⁻⁹, paired two-tailed t-test), as well as an increase of the input resistance (Rin; **Fig. 1 E**) (Control: 87.89 ± 21.79 MΩ [SD]; 4-AP: 108.4 ± 27.80 MΩ [SD]; n = 39 cells, N = 14 mice; p = 3.41 × 10⁻⁸, paired two-tailed t-test) and membrane time constant (Control: 8.79 ± 4.31 ms [SD]; 4-AP: 10.63 ± 5.93 ms [SD]; n = 48 cells, N = 14 mice; p = 0.0159, paired two-tailed t-test; **Fig. 1 F-G**). Altogether, these findings support the notion that I_D_ plays a critical role in shaping both active and passive membrane properties of MesV neurons, consistent with previous work reporting a substantial window current at membrane potentials near the RMP (Dapino et al., 2023).

**Figure 1.**
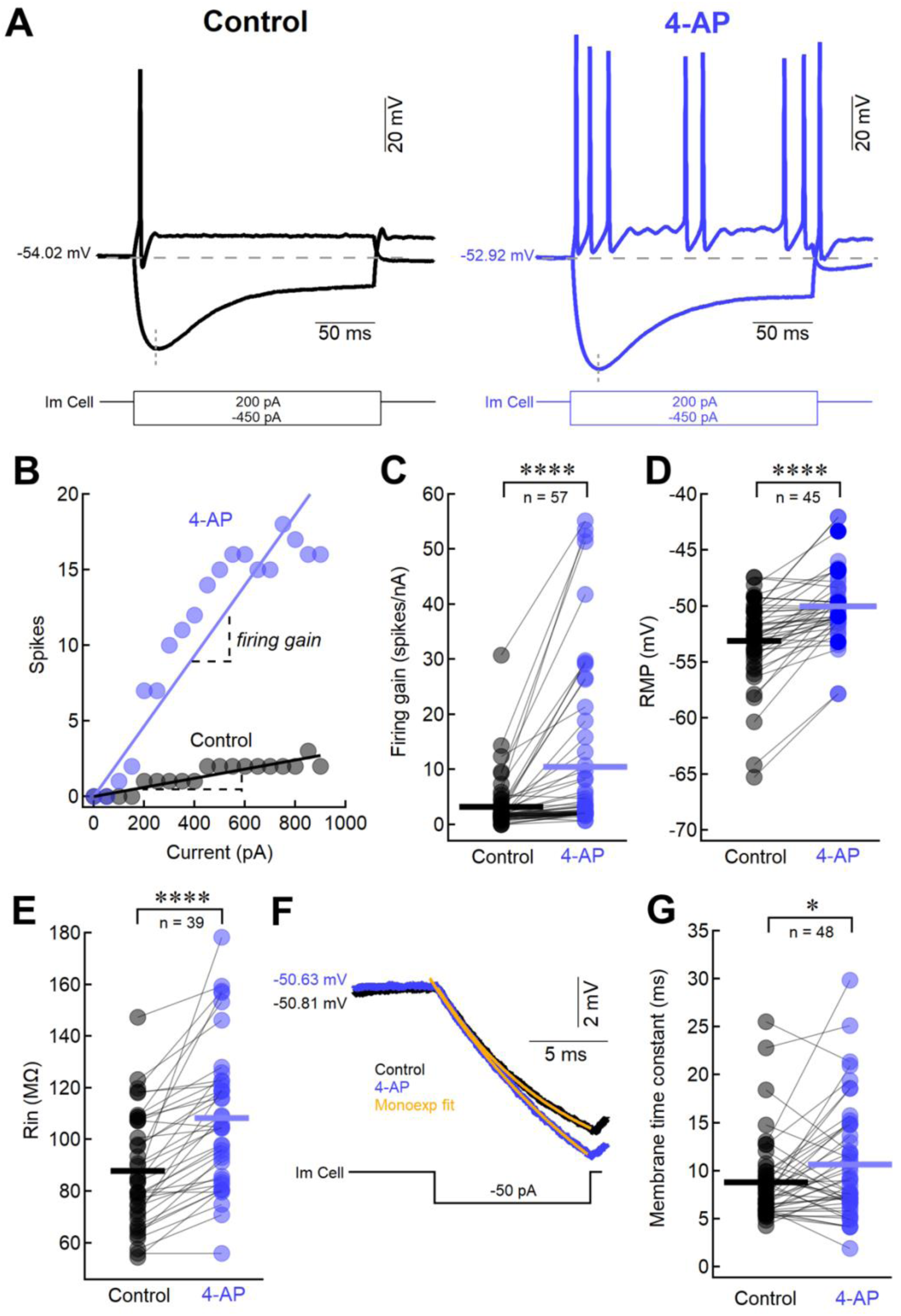
ID contributes to the passive and active electrophysiological properties of MesV neurons. **(A)** Superimposed membrane voltage responses (above) to depolarizing and hyperpolarizing current pulses (below, Im Cell) from a MesV neuron, in control (left) and after bath application of 10 μM 4-AP (right). Values at left of membrane voltage traces denote resting membrane potential in this and subsequent figures. **(B)** Plot of the number of spikes evoked by current pulses of 200 ms in duration as a function of the injected current intensity for the same neuron as in (A), before (black symbols) and after addition of 4-AP (blue symbols). Each data set was fitted to a linear function (superimposed) whose slope represents the neuron’s excitability (firing gain) in each condition. Coupled cells were pulsed independently (unloaded condition, see text). **(C)** Firing gain measured as shown in (B), for the population of recorded MesV neurons in control (black symbols) and in the presence of 4-AP (10 μM; blue symbols) (Control: 3.19 ± 4.67 spikes/nA [SD]; 4-AP: 10.50 ± 15.02 spikes/nA [SD]; n = 57 cells, N = 17 mice; p = 2.27 × 10⁻⁵, paired two-tailed t-test). **(D)** Resting membrane potential (RMP) measured before (black symbols) and after (blue symbols) addition of 10 μM 4-AP (Control: -53.12 ± 3.72 mV [SD]; 4-AP: -50.01 ± 3.13 mV [SD]; n = 45 cells, N = 11 mice; p = 3.85 × 10⁻⁹, paired two-tailed t-test). **(E)** Input resistance (Rin) before (black symbols) and after addition of 4-AP (10 μM; blue symbols) (Control: 87.89 ± 21.79 MΩ [SD]; 4-AP: 108.4 ± 27.80 MΩ [SD]; n = 39 cells, N = 14 mice; p = 3.41 × 10⁻⁸, paired two-tailed t-test). **(F)** Superimposed membrane voltage responses of a MesV neuron (above) to a hyperpolarizing current pulse (below, Im Cell), in control (black trace) and after addition of 10 μM 4-AP (blue trace). These responses were fitted to a single exponential function (orange traces) in order to estimate the membrane time constant (see Materials and Methods). **(G)** Membrane time constant estimated as shown in (F) for the population of recorded MesV neurons before (black symbols) and after addition 4-AP (blue symbols) (Control: 8.79 ± 4.31 ms [SD], 4-AP: 10.63 ± 5.93 ms [SD]; n = 48 cells, N = 14 mice; p = 0.0159, paired two-tailed t-test). Horizontal bars in C, D, E and G, represent population averages.

### The I_D_ of the proximal axon strongly contributes to the somatic electrophysiological properties of MesV neurons

Recent advances have reshaped the classical view of the axon initial segment (AIS), revealing that, beyond acting as the trigger zone for action potential initiation, it also plays a key role in regulating intrinsic neuronal excitability and integration of synaptic inputs (Kole and Stuart, 2012). Consistently, the membrane of the proximal axon contains a rich repertoire of voltage-gated ion channels, in addition to Na⁺ channels. Among these, a high density of voltage-gated K^+^ channels composed of Kv1 subunits has been reported (Dodson et al., 2002; Inda et al., 2006; Shu et al., 2007b; Lorincz and Nusser, 2008; Abdollahi et al., 2025), and these subunits are known contributors to the membrane channels supporting the I_D_ (see below). Based on these antecedents, we investigated whether the proximal axon of MesV neurons expresses the I_D_ and assessed its influence on the electrophysiological properties of the soma, where electrical synapses are located. MesV neurons are pseudounipolar, with a single stem axon that emerges from the soma, that can extend up to several hundred micrometers from the cell body (Baker and Llinás, 1971; Luo et al., 1991). During brainstem slicing, the axons of MesV neurons are transected at varying distances from the soma, resulting in a range of axonal preservation, from complete absence to segments extending several hundred micrometers. During our experiments, the presence of the AIS can be inferred from somatic electrophysiological recordings by assessing the action potential phase plots (dVm/dt vs. Vm), where an abrupt rise (“kink”) is indicative of the AIS-mediated spike (arrow in inset of **Fig. 2 A** and right panel of **Fig. S2 A**) (Bean, 2007; Shu et al., 2007a). Conversely, phase plots lacking this kink suggest that recordings were obtained from neurons devoid of a functional AIS (left panel of **Fig. S2 A**). Therefore, by combining whole-cell electrophysiology with epifluorescence imaging, to visualize the morphology of MesV neurons intracellularly labeled with Alexa Fluor 488 via the recording pipette, we correlated the axon length with the presence or absence of the kink in phase plots. This analysis suggests that a fully functional AIS requires, on average, a proximal axonal length of at least 80 μm (**Fig. 2 A, Fig. S2 A**).

**Figure 2.**
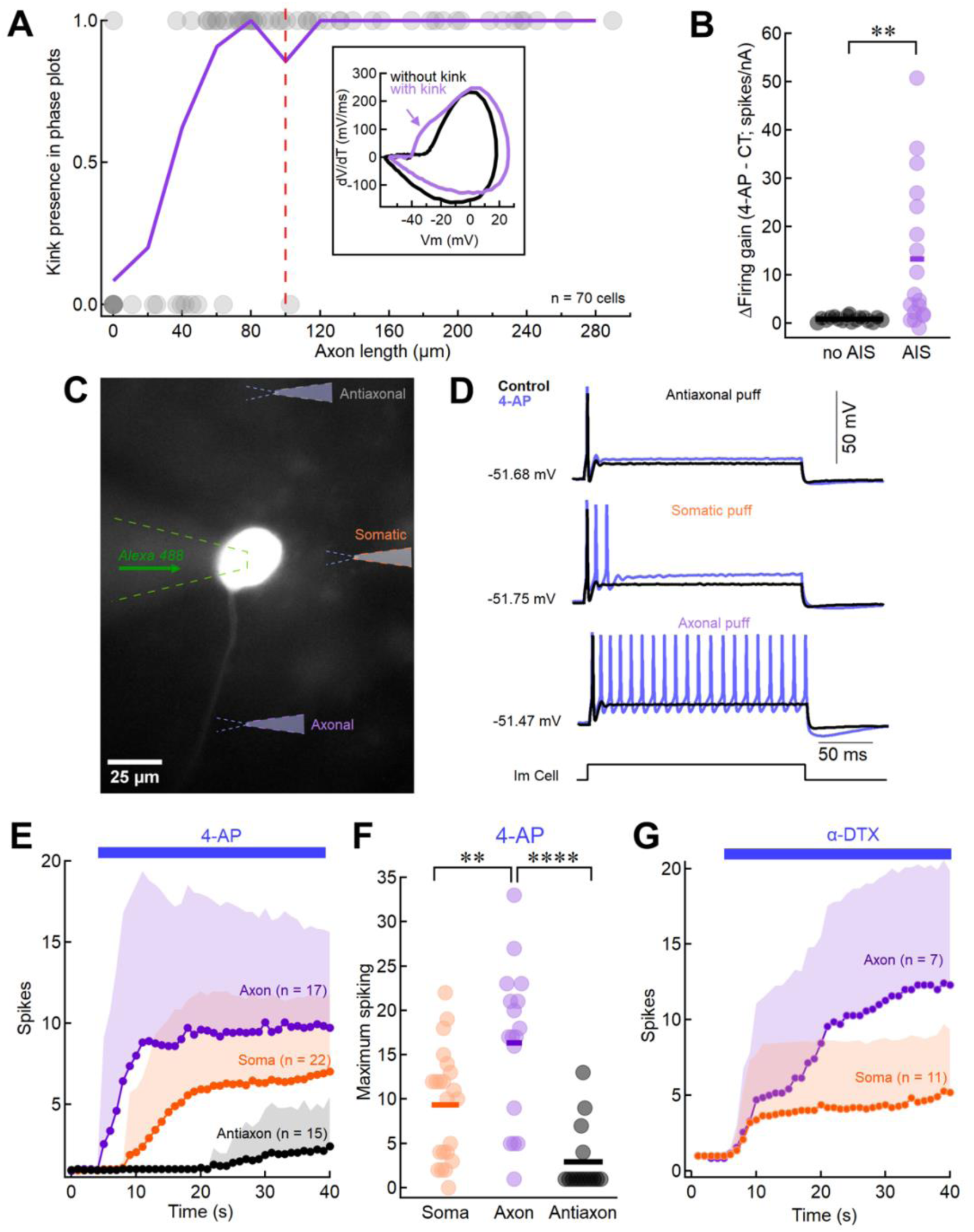
I_D_ of the proximal axon significantly contributes to the somatic excitability of MesV neurons. **(A)** Plot of “kink” incidence in phase plot of recorded neurons (grey symbols; presence = 1, absence = 0), as function of the axonal length measured in MesV neurons intracellularly labeled with Alexa Fluor 488 via the recording pipette. Continuous purple line represents the calculated probability from single cells, binned every 20 μm (n = 70 cells, N = 18 mice). Dashed red line indicates the mean position of the axonal 4-AP puffing pipette (see below). Inset: action potential phase plots (dVm/dt vs. Vm) from two representative MesV neurons are shown superimposed for comparison. One neuron displays a clear kink (purple trace, oblique arrow), indicative of a functional AIS, whereas the other lacks this feature (black trace), suggesting the absence of an AIS. **(B)** Plot of the magnitude of change in the firing gain (ΔFiring gain) before and after 4-AP application for the population of recorded MesV neurons segregated according to absence (black symbols) or presence (purple symbols) of the AIS (No AIS: 0.84 ± 0.46 spikes/nA [SD], n = 17 cells, N = 11 mice; AIS: 13.29 ± 15.11 spikes/nA [SD], n = 18 cells, N = 7 mice; p = 0.0028, unpaired two-tailed t-test with Welch’s correction). **(C)** Picture of a MesV neuron intracellularly labeled with Alexa Fluor 488 via the recording pipette, where the location of the puffing pipettes filled with 4-AP (100 μM) are schematically represented at the axonal, somatic and antiaxonal positions. Scale bar: 25 μm. **(D)** Superimposed membrane voltage responses before (black traces) and during 4-AP injections (blue traces) at the antiaxon (first panel from top), soma (second panel from top) and axon (third panel from top) to a just suprathreshold depolarizing current pulse of 200 ms in duration (fourth trace from top, Im). **(E)** Average plot of spike count per current pulse as a function of time for soma (n = 22 cells, N = 9 mice), axon (n = 17 cells, N = 8 mice) and antiaxon (n = 15 cells, N = 7 mice) 4-AP puffing locations from results like those depicted in (D). Timing of 4-AP (100 μM) application is indicated by the horizontal bar in the upper part of the plot. **(F)** Plot of the maximum spike count per current pulse for each tested neuron during 4-AP (100 μM) puffing applications (Soma: 9.36 ± 6.25 spikes [SD], n = 22 cells, N = 9 mice; Axon: 16.35 ± 8.72 spikes [SD], n = 17 cells, N = 8 mice; Antiaxon: 2.93 ± 3.75 spikes [SD], n = 15 cells, N = 7 mice; axon vs. soma: p = 0.006, unpaired two-tailed t-test; axon vs. antiaxon: p = 7.92 × 10⁻⁶, unpaired two-tailed t-test with Welch’s correction). **(G)** Average plot of spike count per current pulse as a function of time for soma (n = 11 cells) and axon (n = 7 cells) α-DTX puffing locations. Timing of α-DTX (2 μM) application is indicated by the horizontal bar in the upper part of the plot. Shaded area in E and G, represents SD. Horizontal bars in B and F, represent population averages.

Based on the presence of the kink in phase plots, the population of recorded MesV neurons were segregated into two groups: those that retained the AIS, and those in which it was absent. According to the central role the AIS plays in regulating neuronal excitability (Kole and Stuart, 2012), neurons without the AIS showed a significantly more depolarized firing level compared to neurons with it (no AIS: -32.14 ± 4.51 mV [SD], n = 20 cells, N = 13 mice; AIS: -38.29 ± 2.54 mV [SD], n = 46 cells, N = 17 mice; p = 1.35 × 10⁻⁹, unpaired two-tailed t-test; **Fig. S2 B-C**). Also, the Rin is significantly higher in neurons without the AIS compared to neurons with the AIS (no AIS: 146.0 ± 42.52 MΩ [SD], n = 20 cells, N = 13 mice; AIS: 104.9 ± 30.70 MΩ [SD], n = 46 cells, N = 17 mice; p = 5.28 × 10⁻⁵, unpaired two-tailed t-test; **Fig. S2 D-E**), although the RMP did not showed any statistical difference (no AIS: -50.13 ± 1.96 mV [SD], n = 20 cells, N = 13 mice; AIS: -51.40 ± 2.60 mV [SD], n = 46 cells, N = 17 mice; p = 0.0555, unpaired, two-tailed t-test; **Fig. S2 F-G**).

Interestingly, we found that bath applications of 4-AP (10 μM) had a more dramatic effect on MesV neurons that conserved the AIS, compared to those without the AIS. The increase in excitability measured as the magnitude change in firing gain (firing gain in 4- AP minus firing gain in Control, see Material and Methods) averaged 0.84 ± 0.46 spikes/nA [SD] (n = 17 cells, N = 11 mice) in neurons without the AIS, whereas in neurons with the AIS averaged 13.29 ± 15.11 spikes/nA [SD] (n = 18 cells, N = 7 mice; **Fig. 2 B** and **Fig. S2 H-I**). This difference in change of firing gain is statistically significant (p = 0.0028, unpaired two-tailed t-test with Welch’s correction), indicating an increased sensitivity to 4-AP in MesV neurons retaining the AIS. This suggests a high expression of the I_D_ at the AIS of MesV neurons, and that this axonal compartment plays a critical role in shaping somatic electrophysiological properties of these neurons.

To directly assess the expression of the I_D_ at the proximal axon containing the AIS, localized applications (puff) were performed by pressure injections from a second pipette filled with 4-AP (100 μM). Once the recorded neurons were filled with Alexa 488 in order to reveal the presence of the axon and it projecting direction, the puffing pipette was located at the somatic level, the proximal axon (90 μm from the soma on average; range 39-148 μm), or at the anti-axon (same distance as for the axon puff but in the opposite direction), the later as a control for possible diffusional effects to the soma, where the recordings are taken from (**Fig. 2 C**). Regardless of the selected puffing distance, axonal applications were performed exclusively on neurons exhibiting a kink in their action potential phase plot, indicative of a preserved functional AIS. Neuronal excitability was assessed by injecting 200 ms current pulses at 1 Hz, with the intensity adjusted to each neurons rheobase (minimum current for spike activation), both before (5 s) and during puff application. Typically, before puff applications these protocols evoke a single spike at the beginning of current pulses (**Fig. 2 D**, black traces). 4-AP injections promoted an enhancement of neuronal excitability, as evidenced by an increased number of action potentials elicited per current pulse across all three puffing locations (**Fig. 2 D**, blue traces). However, both the time course and the magnitude of this effect varied markedly between sites (**Fig. 2 E and S3 A**-**C**). Indeed, maximum spike firing per pulse, irrespective of its timing of occurrence, averaged 16.35 ± 8.72 spikes [SD] (n = 17 cells, N = 8 mice), 9.36 ± 6.25 spikes [SD] (n = 22 cells, N = 9 mice) and 2.93 ± 3.75 spikes [SD] (n = 15 cells, N = 7 mice), for axon, soma and anti-axon puffing locations respectively. Proximal axon injections had a significative higher effect compared to both, somatic (p = 0.006, unpaired two-tailed t-test) and anti-axon (p = 7.92 × 10⁻⁶, unpaired two-tailed t-test with Welch’s correction) injections (**Fig. 2 F**). Consistent with their effects on spiking, both somatic and proximal axon injections produced a slight but consistent depolarization of the RMP, whereas anti-axon injections had no detectable effect (**Fig. S3 D**). Finally, the possibility that the observed effects were due to a mechanical artifact from puff application, given that MesV neurons express mechanosensitive Piezo2 channels (Florez-Paz et al., 2016), was ruled out by applying vehicle puffs to the soma of identical pressure and duration (**Fig. S3 E**). Vehicle applications produced no changes in membrane excitability (1 ± 0 spikes [SD], n = 9 cells, N = 3 mice, measured at ∼ 33 s from puff onset, corresponding to the time of maximal effect observed with 4-AP: 9.36 ± 6.25 spikes [SD], n = 22 cells, N = 9 mice; p = 4.86 × 10⁻⁶, unpaired, Mann-U Whitney test).

To investigate the ion channel subunits underlying the I_D_, we locally applied α- dendrotoxin (α-DTX, 2 μM) via pressure injections, using the same protocol as in the 4- AP experiments. This toxin selectively blocks channels composed of Kv1.1, Kv1.2, and Kv1.6 subunits, which are known to underlie I_D_ (Bekkers and Delaney, 2001; Haghdoust et al., 2007; Higgs and Spain, 2011). Puff applications of α-DTX at both the soma and proximal axon resulted in an increase of neuronal firing (**Fig. 2 G**), with effects comparable in magnitude to those observed with 4-AP applications (**Fig. S3 F**). Notably, axonal applications evoked a significantly greater increase in spike firing (13.29 ± 7.20 spikes [SD], n = 7 cells, N = 3 mice) compared to somatic applications (6.73 ± 4.92 spikes [SD], n = 11 cells, N = 5 mice; p = 0.035, unpaired two-tailed t-test). These findings strongly suggest that I_D_ is mediated by α-DTX-sensitive Kv1 channels and that these channels are expressed in both somatic and proximal axonal membranes.

The involvement of the proximal axon was further supported by paired whole-cell recordings from the soma and the axonal bleb of the same neuron (**Fig. S4**). Under voltage-clamp conditions, membrane currents were recorded in response to a series of 20 ms hyperpolarizing voltage pulses ranging from 0 to -50 mV in steps of 5 mV, starting from a holding potential of -55 mV, which were alternately applied to each compartment (**Fig. S4 A**). From these recordings, plots of the membrane current in the non-stepped compartment (which corresponds to the current flowing between compartments) as a function of the axon-soma voltage difference were constructed, and the intercompartment conductance determined from the slope of linear regressions (**Fig. S4 B**). As expected, the conductance values determined in this way were nearly symmetrical in opposite directions (average difference expressed as percentage of the soma-axon direction: 4.4%, **Fig. S4 C**). From these recordings, the coupling coefficient was estimated for each direction (Curti and O’Brien, 2016) yielding values that ranged from 0.3 to 0.6, (n = 3 cells, N = 3 mice; **Fig. S4 D**). These findings indicate a relatively strong electrotonic coupling between the two compartments, supporting the notion of bidirectional interactions between the soma and proximal axon in MesV neurons. In summary, these results confirm a substantial expression of the I_D_ in the membrane of the proximal axon and highlight the critical contribution of this compartment to the somatic electrophysiological properties, thus impacting on electrical synaptic transmission between MesV neurons.

### The I_D_ contributes to the coupling strength between MesV neurons

Preceding results showing the contribution of I_D_ to the passive membrane properties of MesV neurons, particularly influencing Rin (**Fig. 1 E**, see above), suggest that it may also participate in setting the coupling strength between these neurons (Bennett, 1966; Getting, 1973). To test this hypothesis, we performed simultaneous whole-cell recordings from pairs of MesV neurons. Electrical coupling was confirmed in current clamp, by injecting a series of hyperpolarizing current steps (0 to -450 pA) into one neuron while membrane potential changes were recorded in both cells (**Fig. 3 A**). From these recordings, the coupling coefficient (CC) was calculated (see Materials and Methods) under control conditions and following application of 4-AP (10 μM). Partial blockade of the I_D_ (∼75%) led to a significant increase in CC (Control: 0.27 ± 0.17 [SD]; 4-AP: 0.31 ± 0.17 [SD]; n = 45 directions, N = 14 mice; p = 0.0089, paired two-tailed t-test; **Fig. 3 B-C**). These results highlight a relevant role of the I_D_ in modulating the strength of electrical transmission for hyperpolarizing signals, most probably due to its steady state activation at membrane potentials around the RMP (Dapino et al., 2023). Despite that, many network operations supported by electrical synapses such as synchronization and coincidence detection, primarily rely on the transmission of depolarizing potentials. To directly evaluate whether the I_D_ also contributes to the efficacy of electrical transmission in the depolarizing range, we quantified the spike-related coupling coefficient (CC_spike_), based on recordings such as those shown in **Figure 3 D**. The CC_spike_ increased significantly following partial I_D_ blockade with 4-AP, rising from 0.055 ± 0.023 [SD] in control conditions to 0.074 ± 0.026 [SD] after 4-AP (n = 16 directions, N = 7 mice; p = 0.0001, paired two-tailed t-test; **Fig. 3 E**). Notably, while partial blockade of the I_D_ led to an increase in the CC of approximately 15%, the CC_spike_ increased by about 35%. The more pronounced effect of 4-AP on CC_spike_ suggests that, whereas changes in CC likely reflect the contribution of the I_D_ to Rin, the enhancement of CC_spike_ reveals an additional role of the I_D_ in attenuating the amplitude of spike-evoked coupling potentials. Together, these findings indicate that I_D_ contributes to the regulation of electrical coupling strength between MesV neurons, both by influencing postsynaptic input resistance and by dynamically opposing depolarizing coupling potentials such as spikelets, as suggested previously (Dapino et al., 2023). On the other hand, the latency of transmission, measured as the peak-to-peak interval between the presynaptic spike and the postsynaptic spikelet, increased from 0.43 ± 0.13 ms [SD] in control conditions to 0.96 ± 0.45 ms [SD] after 4-AP (10 μM) (n = 16 directions, N = 7 mice; p = 0.00002, paired two-tailed t-test; **Fig. 3 F**). This suggests that I_D_ activation shortens the time-to-peak of the spikelet, thereby also contributing to the temporal characteristics of transmission.

**Figure 3.**
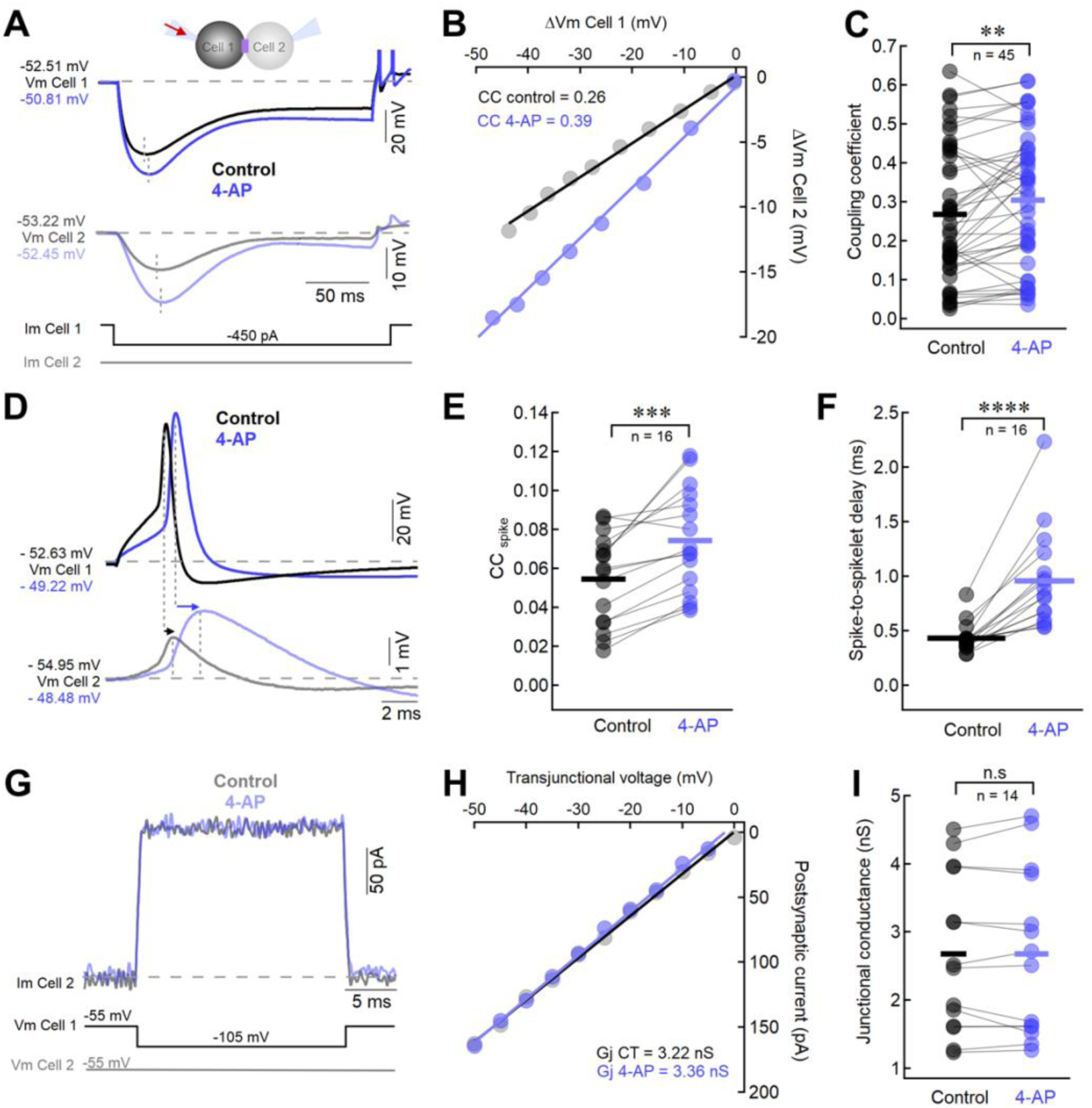
I_D_ determines the strength and temporal characteristics of electrical synaptic transmission between MesV neurons. **(A)** Paired recordings from a pair of electrically coupled MesV neurons in response to a hyperpolarizing current pulse (Im Cell 1) showing the membrane voltage response in the injected (Vm Cell 1) and coupled (Vm Cell 2) neurons, in control (black traces) and in the presence of 4-AP (10 μM) (blue traces). The scheme above traces indicates the experiment configuration. **(B)** From recordings like those depicted in (A) in which a series of hyperpolarizing current pulses of increasing intensity were injected in one neuron, the coupling coefficient (CC) was estimated by plotting the amplitude of membrane voltage changes (measured at the peak of hyperpolarizing responses, vertical dashed lines in A) in the postsynaptic cell (ΔVm Cell 2) as a function of membrane voltage changes in the presynaptic cell (ΔVm Cell 1), in control (black symbols) and in the presence of 4-AP (blue symbols). Each data set was fitted with a linear function whose slope values representing the CC are indicated. **(C)** Plot of CC values measured in control (black symbols) and after the addition of 10 μM 4-AP (blue symbols) for the population of assessed directions (Control: 0.27 ± 0.17 [SD]; 4-AP: 0.31 ± 0.17 [SD]; n = 45 cells, N = 14 mice; p = 0.0089, paired, two-tailed t-test). **(D)** Spike transmission, showing the presynaptic spike (above, Vm Cell 1) elicited by a short depolarizing current pulse, and the corresponding coupling potential in the postsynaptic neuron (below, Vm Cell 2), in control (black traces) and after 4-AP (blue traces) application. Arrows denote presynaptic spike to postsynaptic spikelet transmission delay. **(E)** Spike-related coupling coefficient (CC spike) for the population of assessed directions, calculated from recordings like those depicted in (D) in control and after 10 μM 4-AP (Control: 0.055 ± 0.023 [SD]; 4-AP: 0.074 ± 0.026 [SD]; n = 16 directions, N = 7 mice; p = 0.0001, paired, two-tailed t-test). **(F)** Spike-to-spikelet delay transmission measured as depicted in (D), in control and after 4-AP (10 μM) (Control: 0.43 ± 0.13 ms [SD]; 4-AP: 0.96 ± 0.45 ms [SD]; n = 16 directions, N = 7 mice; p = 0.00002, paired, two-tailed t-test). **(G)** Voltage clamp recording from a pair of electrically coupled MesV neurons during the application of a hyperpolarizing step-like voltage command in one cell (Vm Cell 1) while the membrane potential of the other cell was held constant (Vm Cell 2), showing the membrane current response elicited in the non-stepped neuron (above, Im Cell 2), corresponding to the junctional current in control (black trace) and in the presence of 4-AP (30 μM) (blue trace). **(H)** From recordings like those depicted in (G) in which a series of hyperpolarizing voltage commands of increasing magnitude were applied in one neuron, the junctional conductance (Gj) was estimated by plotting the amplitude of the postsynaptic membrane current as a function of the membrane voltage difference between cells (transjunctional voltage), in control (black symbols) and in the presence of 4-AP (blue symbols). Each data set was fitted with a linear function and the slope values representing the Gj are indicated. **(I)** Gj values for the population of assessed direction in control and after 4-AP (30 μM) (Control: 2.68 ± 1.16 nS [SD]; 4-AP: 2.68 ± 1.22 nS [SD]; n = 14 directions from 7 pairs, N = 3 mice; p = 0.9500, paired, two-tailed t-test). Horizontal bars in C, E, F and I represent population averages.

Importantly, the increase in CC and CC_spike_ were not accompanied by changes in junctional conductance (Gj), as confirmed by paired voltage-clamp recordings from electrically coupled MesV neurons (**Fig. 3 G**; see Materials and Methods). Gj values were similar under control conditions and after 4-AP application (30 μM) (Control: 2.68 ± 1.16 nS [SD]; 4-AP: 2.68 ± 1.22 nS [SD]; n = 14 directions, N = 3 mice; p = 0.9500, paired two-tailed t-test; **Fig. 3 H-I**). This highlights the contribution of mechanisms of the non-junctional membrane, particularly the I_D_, to the coupling strength.

### The I_D_ regulates the gain of coincidence detection

In previous work, we demonstrated that electrical coupling between MesV neurons in rats supports coincidence detection, and that intrinsic electrophysiological properties can significantly regulate this phenomenon (Curti et al., 2012; Davoine and Curti, 2019). Based on our current findings indicating that I_D_ contributes to both spiking properties and coupling strength (**Figs. 1** and **3**), we next examined whether this conductance contributes to coincidence detection.

To assess this, we first evaluated the ability of electrically coupled MesV neurons in mice to detect coincident (simultaneous) inputs using established protocols (Curti et al., 2012). During paired recordings of electrically coupled neurons, injection of a just subthreshold current pulse into one neuron elicited a local voltage response, and a corresponding coupling potential in the other neuron (**Fig. 4 A**, left panel). In contrast, when current pulses of the same intensity were delivered simultaneously to both neurons, action potentials were triggered in both cells (**Fig. 4 A**, right panel), indicating that MesV neurons are more responsive to coincident depolarizing inputs. This finding was confirmed by the quantification of the rheobase, showing significantly lower values during independent versus simultaneous stimulation (Independent: 362.5 ± 146.2 pA [SD]; Simultaneous: 328.6 ± 133.4 pA [SD]; n = 56 cells, N = 16 mice; p = 3.43 × 10⁻⁷, paired two-tailed t-test; **Fig. 4 B**). Altogether, these results indicate that electrically coupled MesV neurons in mice can operate as coincidence detectors.

**Figure 4.**
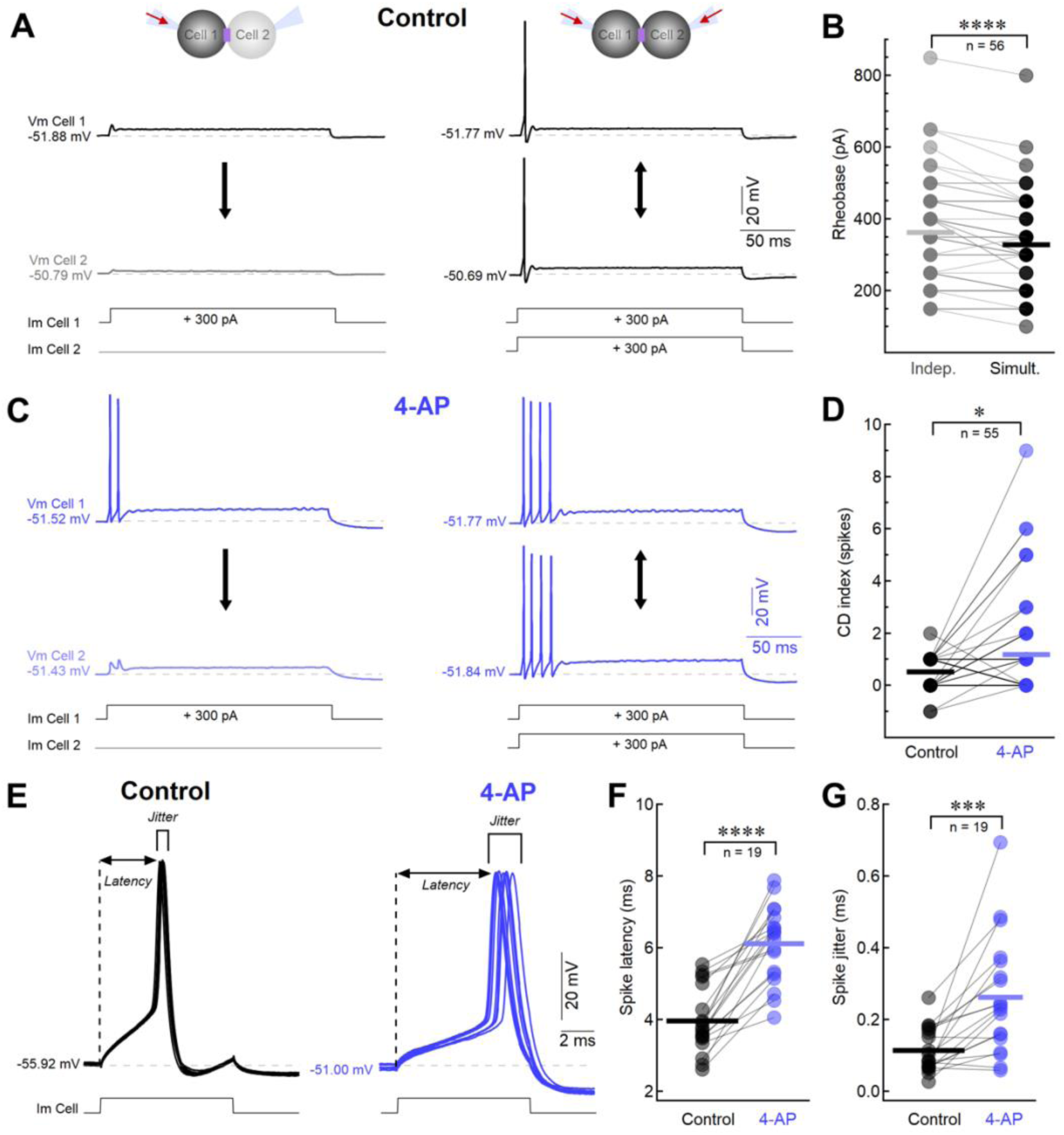
I_D_ of MesV neurons contributes to set the gain of coincidence detection. **(A)** Membrane voltage responses of a pair of coupled MesV neurons (Vm Cell 1 and Vm Cell 2) to a current pulse delivered only to Cell 1 (left), and to both cells at the same time (Im Cell 1 and Im Cell 2) (right). The schemes above each set of traces indicate the experiment configuration. **(B)** Minimum intensity for spike activation of a 200 ms current pulse (Rheobase) when independently (Indep.) or simultaneously (Simult.) applied from recordings like those shown in (A) (Independent: 362.5 ± 146.2 pA [SD]; Simultaneous: 328.6 ± 133.4 pA [SD]; n = 56 cells, 28 pairs, N = 16 mice; p = 3.43 × 10⁻⁷, paired two-tailed t-test). **(C)** For the same pair of MesV neurons depicted in (A), membrane voltage responses to the same stimulating protocol applied in the presence of 4-AP (10 μM). **(D)** Coincidence detection (CD) index, defined as the difference in spike count when stimuli were delivered simultaneously versus individually in control and after the addition of 4 - AP (Control: 0.52 ± 0.60 spikes [SD]; 4-AP: 1.18 ± 1.96 spikes [SD]; n = 55 cells, 28 pairs, N = 16 mice; p = 0.0182, paired two-tailed t-test). **(E)** Superimposed membrane voltage responses of a MesV neuron to 10 consecutives just-suprathreshold current pulses (Im Cell) delivered at 0.5 Hz, in control (left) and after the application of 10 μM 4-AP (right). From recordings such as those shown in (E), spike latency **(F)** and spike jitter **(G)** were quantified across the recorded neuronal population. Spike latency was defined as the time from current pulse onset to the spike peak (horizontal double arrows in E) (Control: 3.97 ± 0.90 ms [SD]; 4-AP: 6.11 ± 1.05 ms [SD]; n = 19 cells, 10 pairs, N = 5 mice; p = 1.00 × 10⁻⁷, paired two-tailed t-test), and spike jitter as the corresponding standard deviation (Control: 0.11 ± 0.06 ms [SD]; 4-AP: 0.26 ± 0.16 ms [SD]; n = 19 cells, 10 pairs, N = 5 mice; p = 0.0004, paired two-tailed t-test). Horizontal bars in B, D, F and G represent population averages.

To determine the role of the I_D_ in this phenomenon, we repeated the stimulation protocol in the presence of 4-AP (10 μM). Under these conditions, independent current pulses that previously failed to elicit spiking, now generated action potentials (**Fig. 4 C**, left panel), consistent with increased neuronal excitability following partial blockade of the I_D_ (**Fig. 1**). Furthermore, simultaneous current injections induced robust, repetitive firing in both neurons (**Fig. 4 C**, right panel). The gain of coincidence detection was quantified using a coincidence detection index (CD index), defined as the difference in spike count when stimuli were delivered simultaneously versus individually (Davoine and Curti, 2019). The CD index increased significantly from 0.52 ± 0.60 spikes [SD] under control conditions to 1.18 ± 1.96 spikes [SD] after partial I_D_ blockade (n = 55 cells, N = 16 mice; p = 0.0182, paired two-tailed t-test; **Fig. 4 D**). Consistently, the reduction of the rheobase during coincident stimulation respect to independent stimulation, was significantly larger after 4-AP (10 μM) application (reduction in control: 33.93 ± 43.80 pA [SD]; reduction in 4-AP: 61.61 ± 54.76 pA [SD]; n = 56 cells, 28 pairs, N = 16 mice; p = 9.91× 10⁻⁶, paired two-tailed t-test; data not shown).

Moreover, the I_D_ also contributes to the temporal characteristics of firing due to coincident inputs (**Fig. 4 E**). In fact, during simultaneous activation of pairs of coupled MesV neurons, spike latency increased significantly from 3.97 ± 0.90 ms [SD] under control conditions, to 6.11 ± 1.05 ms [SD] following application of 4-AP (10 μM) (n = 19 cells, N = 5 mice; p = 1.00 × 10⁻⁷, paired two-tailed t-test; **Fig. 4 F**). Similarly, spike jitter (measured as the standard deviation of spike latency across repeated stimulations) increased from 0.11 ± 0.06 ms [SD] in control, to 0.26 ± 0.16 ms [SD] in 4-AP (n = 19 cells, N = 5 mice; p = 0.0004, paired two-tailed t-test; **Fig. 4 G**). Altogether, these results indicate that by dampening the neuronal excitability, the I_D_ sets the gain of coincidence detection, as well as the timing of firing during simultaneous depolarizing inputs to electrically coupled neurons.

### The I_D_ shapes the precision of coincidence detection

The contribution of the I_D_ to set the temporal characteristics of electrical transmission between MesV neurons (**Fig. 3 D** and **F**) suggests that this conductance may also contribute to establish the precision of coincidence detection. To confirm this hypothesis, we quantified the time window over which pairs of electrically coupled MesV neurons exhibit coincidence detection. This was done by delivering near-threshold, short-duration (3 ms) current pulses to each neuron with systematically varied inter-pulse delays in 0.5 ms increments (Alcami, 2018) (**Fig. 5 A**) and measuring changes in firing probability. Spike counts from both neurons were summed and plotted as a function of inter-stimuli delay. As expected, in control conditions, spike probability was maximal at 0 ms delay (synchronous input) and declined progressively with increasing delays (**Fig. 5 B**, black symbols). Population-averaged data exhibited a bell-shaped relationship that was well fitted to a Gaussian function (**Fig. 5 C**, black symbols). Notably, partial blockade of the I_D_ markedly broadened this temporal window (**Fig. 5 B**, blue symbols). The Gaussian fit width representing the time interval between stimuli to give the half-maximum firing probability (full width at half maximum, FWHM) more than doubled, increasing from 4.01 ± 0.12 ms [SD] in control conditions to 10.25 ± 0.43 ms [SD] in 4-AP (10 μM) (**Fig. 5 C**, blue symbols). Consistently, statistical analysis revealed that the distributions in control and in 4-AP are statistically different (p = 8.8 × 10⁻^35^, Kolmogorov–Smirnov test). Altogether, these findings indicate that the I_D_ plays a critical role in defining the temporal properties of coincidence detection mediated by electrical synapses, by enhancing its temporal precision at the cost of reduced gain.

**Figure 5.**
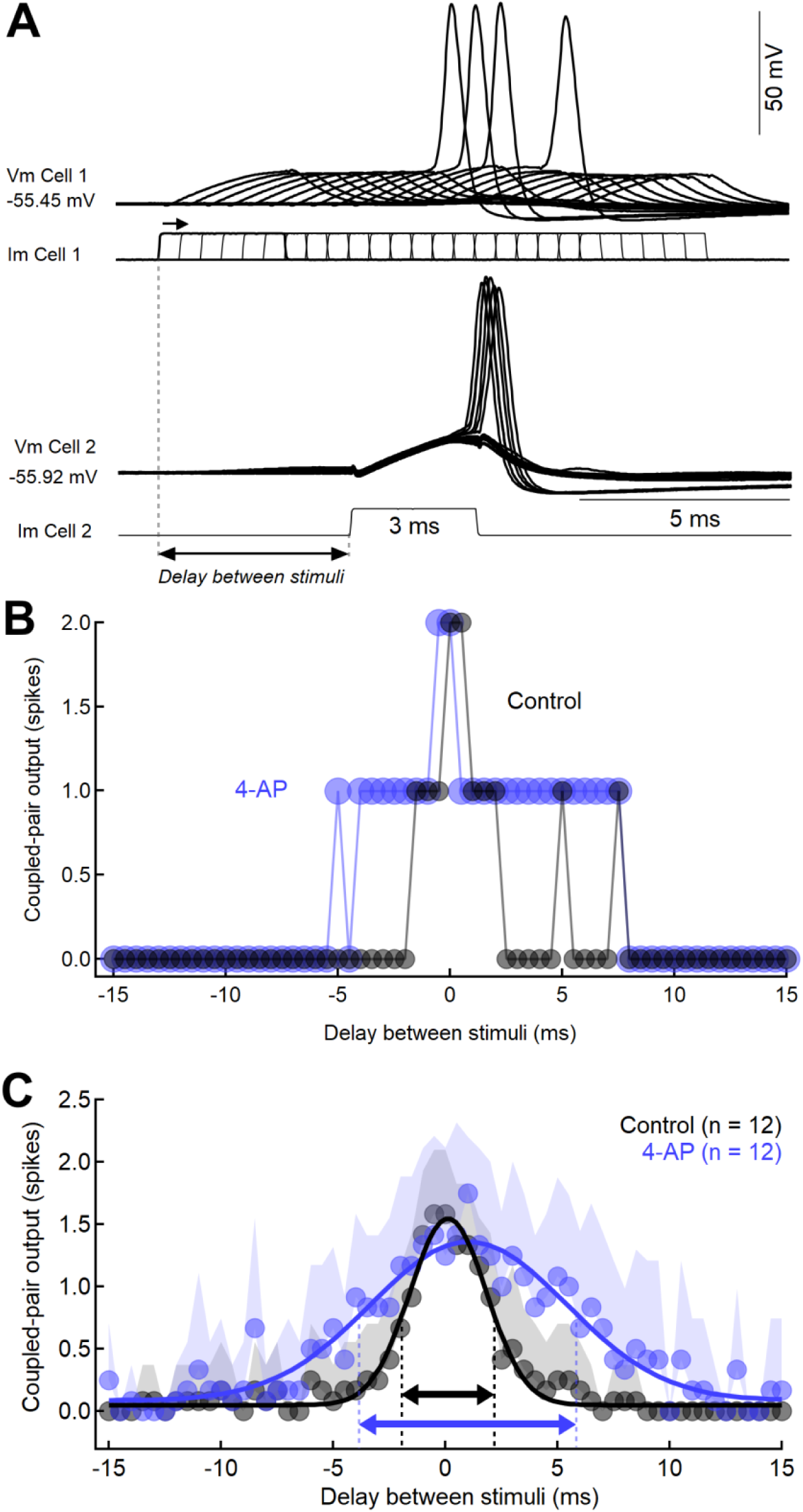
I_D_ of MesV neurons contributes to set the precision of coincidence detection. **(A)** Superimposed membrane voltage responses of a pair of coupled MesV neurons (Vm Cell 1 and Vm Cell 2) to a stimulation protocol consisting of near-threshold, short-duration (3 ms) current pulses delivered to each neuron (Im Cell 1 and Im Cell 2). The timing of the current pulses in one neuron (Cell 2, in this example) was kept constant, while in the other neuron (Cell 1) it was systematically varied in 0.5 ms increments. **(B)** For the pair of coupled MesV neurons shown in (A) spike counts from both neurons were summed and plotted as a function of inter-stimuli delay in control (black symbols) and in the presence of 10 μM 4-AP (blue symbols). **(C)** Average plot of spike counts from both neurons (Coupled-pair output) as a function of inter-stimuli delay for the population of recorded MesV neurons coupled pairs in control (black symbols) and in the presence of 10 μM 4-AP (blue symbols). Shaded area represents SD. Superimposed are Gaussian fits to the data. Double arrows indicate the full width at half maximum (FWHM) of each fit, with values of 4.01 ms for control (black) and 10.25 ms for 4-AP (blue). Gaussian fit parameters in control: y0 = 0.05 ± 0.008 spikes [SD], Amplitude = 1.5 ± 0.04 spikes [SD], x0 = 0.12 ± 0.05 ms [SD], width = 2.41 ± 0.07 ms [SD], FWHM = 4.01 ± 0.12 ms [SD]; in 4-AP: y0 = 0.085 ± 0.017 spikes [SD], Amplitude = 1.27 ± 0.0425 spikes [SD], x0 = 1.03 ± 0.16 ms [SD], width = 6.15 ± 0.26 ms [SD], FWHM = 10.25 ± 0.43 ms [SD]; n = 12 coupled pairs, N = 8 mice; paired representation; paired distribution comparison: p = 8.8 × 10⁻^35^, Kolmogorov–Smirnov test).

## DISCUSSION

In this study, we investigated the role of the I_D_, a subthreshold voltage-gated K⁺ conductance, in coincidence detection within circuits of electrically coupled neurons. Coincidence detection is a circuit property whereby input timing differences are encoded in the firing rates of postsynaptic neurons, enabling preferential responses to synchronous inputs within the microsecond-to-millisecond range, compared to temporally dispersed ones. This mechanism is critical for sensory processes such as sound source localization, where it has been extensively studied in binaural neurons of the auditory system in birds (Carr and Konishi, 1990; Joseph and Hyson, 1993; Overholt et al., 1992) and mammals (Spitzer and Semple, 1995; Yin and Chan, 1990). Morphologically, these neurons —particularly those tuned to low and medium frequencies— have bipolar dendrites, each receiving input from one ear (Joris et al., 1998). This anatomical segregation renders inputs to a single dendrite relatively ineffective for spike initiation, as part of the incoming current is shunted to the contralateral dendrite, a phenomenon analogous to the loading effect previously described (see Introduction). In contrast, coincident inputs to both dendrites mitigate this shunting because voltage changes occur simultaneously in both sides, leading to strong depolarization of the soma and proximal axon, supporting reliable action potential generation. Thus, each dendrite effectively acts as a current sink for inputs to the other, enabling coincidence detection (Agmon-Snir et al., 1998). This mechanism closely parallels coincidence detection in circuits of electrically coupled neurons (Bennett and Zukin, 2004; Curti et al., 2022). In fact, in pairs of coupled MesV neurons, each cell serves as a current sink for its partner when only one neuron undergoes depolarization. However, synchronous inputs to both neurons minimize voltage differences across gap junctions, reducing current shunting and increasing the likelihood of firing. While the coincidence detection window differs between systems (∼4 ms in MesV neurons vs. ∼2 ms at 20 °C in nucleus laminaris neurons; Kuba, 2003), the underlying principle remains the same: electrotonically coupled compartments receiving segregated inputs. In the auditory system, this phenomenon, combined with specialized circuit arrangements such as delay lines, supports computation of interaural time differences for sound localization (Joris et al., 1998; Kuba, 2003). Notably, MesV neurons share several properties with coincidence detector neurons of the nucleus laminaris and the medial superior olive. They exhibit relatively fast membrane time constants and generate a single spike at the onset of suprathreshold current pulses, reflecting their phasic firing behavior (Kuba et al., 2005; Scott et al., 2005). In addition, MesV neurons display pronounced outward and inward rectification near resting membrane potential, mediated by low-threshold K⁺ currents and hyperpolarization-activated cationic currents, respectively (Del Negro and Chandler, 1997; Khakh and Henderson, 1998; Tanaka et al., 2003), closely resembling the properties of auditory system neurons (Kuba et al., 2002). The functional relevance of coincidence detection in orofacial behaviors involving MesV neurons, however, remains unclear. These primary afferents, located in the brainstem, receive extensive synaptic input from diverse regions via multiple neurotransmitter systems, suggesting they also function as interneurons within circuits controlling jaw movements (Lazarov and Chouchkov, 1995; Lazarov and Pilgrim, 1997; Lazarov, 2002; Liem et al., 1997; Verdier et al., 2004; Takahashi et al., 2010; Fortin et al., 2021). In this context, electrical coupling likely enables MesV neurons to preferentially respond to temporally clustered excitatory inputs from hierarchically superior structures, as opposed to temporally dispersed inputs.

Our findings demonstrate that the electrophysiological properties of MesV neurons are strongly shaped by the I_D_, consistent with our previous work (Dapino et al., 2023). This current is expressed both at the soma and proximal axon, likely mediated by Kv1- containing channels. Notably, these compartments are tightly electrotonically coupled, enabling bidirectional interactions. This supports the idea that voltage-dependent mechanisms at the AIS significantly influence somatic properties, including those relevant to electrical synaptic transmission. The I_D_ contributes to the coupling strength of depolarizing signals in two ways. First, its large window current, resulting from overlap between activation and inactivation curves (Dapino et al., 2023), provides steady-state activation near resting potential. This lowers Rin, and consequently, coupling strength. Second, the rapid activation kinetics of the I_D_ act as a dynamic brake on depolarizations, such as spike-induced coupling potentials (spikelets). This not only dampens coupling efficacy but also shortens the duration of spikelets by accelerating their falling phase. This latter effect reduces spikelet time-to-peak and minimize presynaptic-to-postsynaptic delays. Together, these properties allow the I_D_ to finely regulate both the strength and temporal precision of electrical transmission.

As time management is essential for coincidence detection, any membrane mechanism that defines a neuron’s integration window—the interval during which synaptic inputs are effectively summed—directly influences its temporal precision. In this context, the activation range and rapid kinetics of the I_D_ make it particularly well-suited to shape excitatory synaptic potentials, spike timing, and action potential repolarization. Our results support this role, showing that the I_D_ significantly contributes to the precision of coincidence detection, consistent with prior experimental findings in nucleus laminaris neurons (Kuba et al., 2002; Kuba, 2003) and modeling results in the medial superior olive (Mathews et al., 2010). However, while the I_D_ sharpens voltage changes over time, it simultaneously dampens membrane excitability. This dual action reduces the contrast between minimal responses to asynchronous inputs and maximal responses to synchronous ones. In summary, the I_D_ critically modulates coincidence detection by enhancing its temporal precision while limiting its gain.

## Acknowledgements

Funding: this work was supported by Agencia Nacional de Investigación e Innovación (ANII), Uruguay (FCE_1_2021_1_166745); Programa de Desarrollo de las Ciencias Básicas (PEDECIBA); and Comisión Académica de Posgrado of Universidad de la República, Uruguay.

---

**Supplementary Figure 1.**
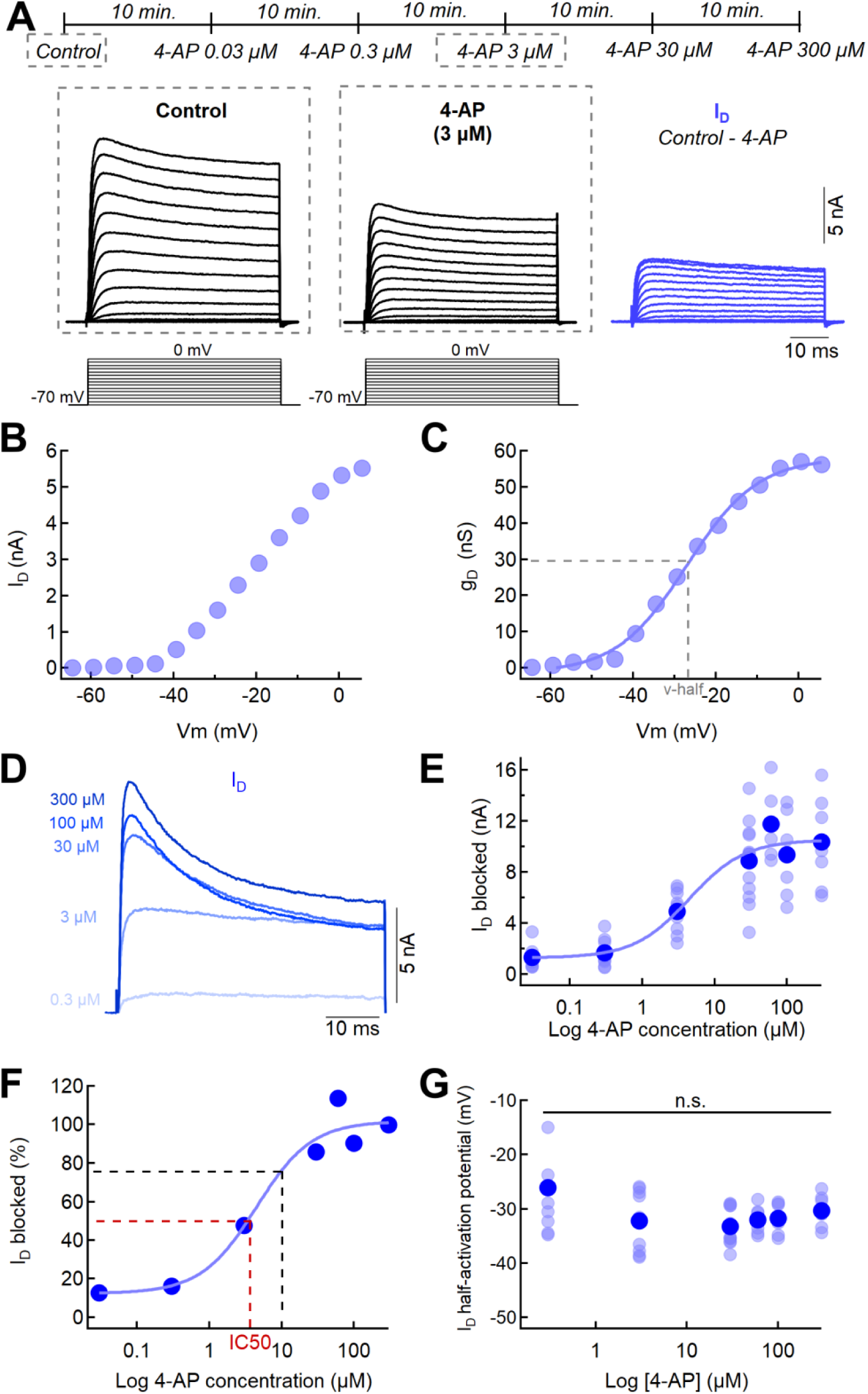
Characterization of I_D_ sensitivity to 4-AP in MesV neurons. **(A)** *Top*: timeline of experimental procedure in which recorded cells were exposed to increasing extracellular concentrations of 4-AP (0.03, 0.3, 3, 30 and 300 μM) every 10 min. starting from the control condition (see Materials and Methods). *Middle:* representative membrane current recordings in response to a series of 50 ms voltage steps from 0 to 70 mV in increments of 5 mV, starting from a holding potential of -70 mV (bottom) control (first panel from left) and 4-AP (3 μM, second panel from left). To isolate the I_D_, current recordings in 4-AP were subtracted from those obtained in control conditions (third panel from left). **(B)** Plot of the maximum I_D_ as a function of membrane voltage from the recordings shown in (A). **(C)** Activation curve of the I_D_ constructed from the data shown in (B). Fits to a Boltzmann function (continuous trace) is superimposed to the experimental data (round symbols) and the vertical dashed line indicate the half-activation voltage value. **(D)** Representative recordings of the isolated I_D_ obtained employing different concentrations of 4-AP in the same neuron. **(E)** Magnitude of the ID I_D_ blocked as a function of the 4-AP concentration for the population of recorded neurons. Superimposed are the individual values (light blue circles) and the corresponding averages (dark blue circles; 0.3 μM: 1.67 ± 1.09 nA [SD], n = 9 cells, N = 5 mice; 3 μM: 4.93 ± 1.64 nA [SD], n = 9 cells, N = 5 mice; 30 μM: 8.88 ± 3.02 nA [SD], n = 13 cells, N = 6 mice; 60 μM: 11.75 ± 2.75 nA [SD], n = 6 cells, N = 3 mice; 100 μM: 9.35 ± 3.19 nA [SD], n = 7 cells, N = 3 mice; 300 μM: 10.34 ± 3.56 nA [SD], n = 7 cells, N = 3 mice). Also superimposed is the fit to a Hill equation. **(F)** Percentage of I_D_ blockade as a function of 4-AP concentration obtained from data shown in (E). Superimposed is the fit to a Hill equation (base: 12.42 ± 10.2 %; max: 101.84 ± 11.3 %; rate: 1.13 ± 0.83 %; xhalf [IC50]: 4.45 ± 2.69 μM). Dashed vertical lines indicate the value of the IC50 (red) and the concentration used to evaluate the contribution of the I_D_ to the excitability, coupling strength and coincidence detection in the present study (black). **(G)** Half activation voltage values of the I_D_ at each 4-AP concentration employed determined as shown in (C). Superimposed are the individual values (light blue circles) and the corresponding averages (dark blue circles; 0.3 μM: -26.05 ± 9.04 mV [SD], n = 9 cells, N = 5 mice; 3 μM: -32.24 ± 5.72 mV [SD], n = 9 cells, N = 5 mice; 30 μM: -33.24 ± 3.18 mV [SD], n = 13 cells, N = 6 mice; 60 μM: -32.06 ± 2.63 mV [SD], n = 6 cells, N = 3 mice; 100 μM: -31.72 ± 2.81 mV [SD], n = 7 cells, N = 3 mice; 300 μM: -30.33 ± 2.95 mV [SD], n = 7 cell, N = 3 mice; p = 0.0512, one-way ANOVA).

**Supplementary Figure 2.**
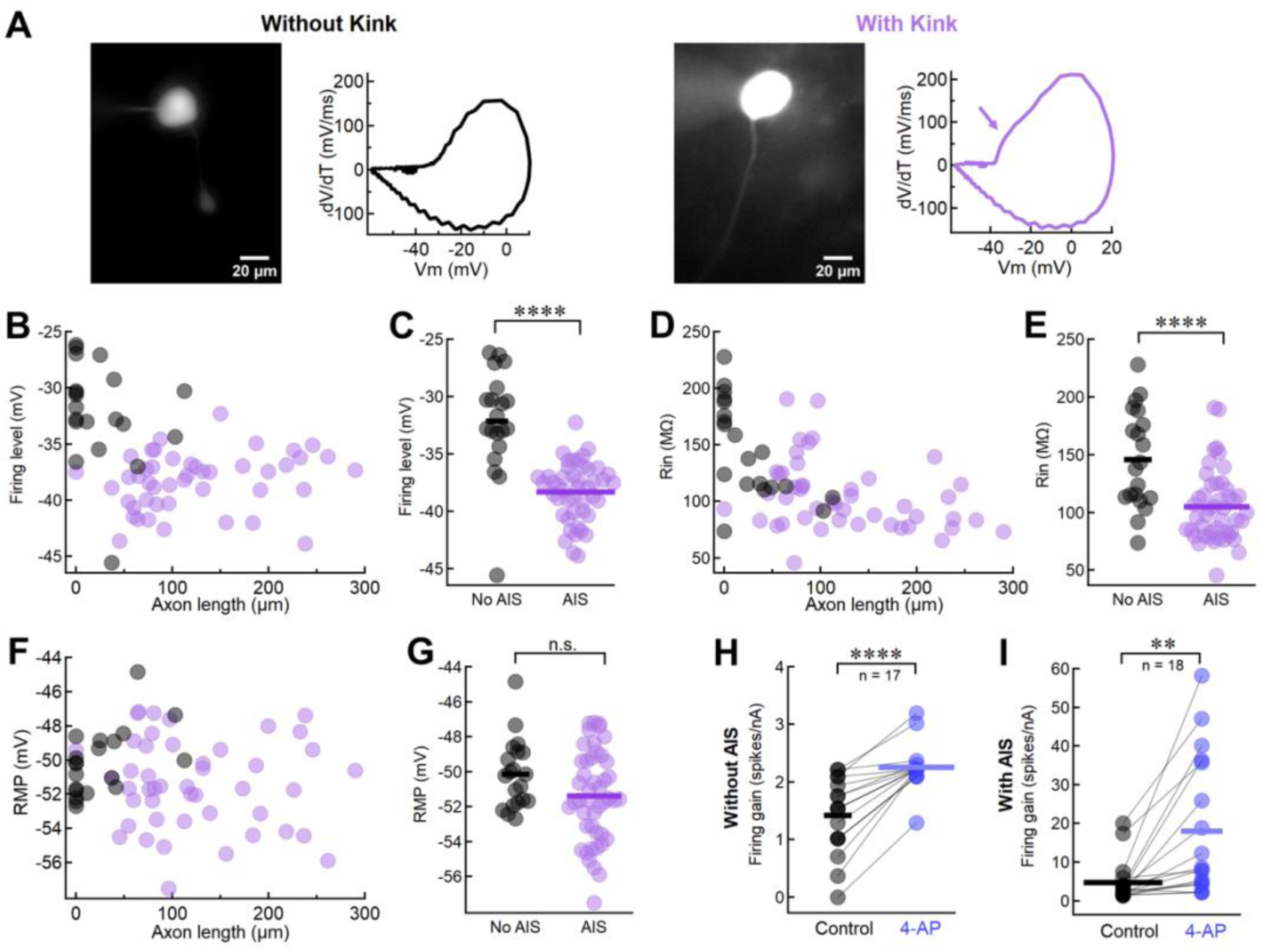
Contribution of the proximal axon membrane to the somatic electrophysiological properties of MesV neurons. **(A)** Representative images of two MesV neurons intracellularly labeled with Alexa Fluor 488 via the recording pipette, shown alongside their corresponding phase plots. The neuron on the right lacks a kink, indicative of AIS absence, whereas the neuron on the left displays a kink (oblique arrow), indicative of AIS presence. **(B)** Firing level as a function of axon length for the population of recorded neurons (n = 66 cells, N = 18 mice), determined from experiments shown in (A). The plot displays values from neurons with an AIS (purple symbols) and without an AIS (grey symbols), according to the presence or absence of a kink in their phase plots. **(C)** Firing level values of neurons with AIS and without AIS from the same data displayed in (B) (No AIS: -32.14 ± 4.51 mV [SD], n = 20 cells, N = 13 mice; AIS: -38.29 ± 2.54 mV [SD], n = 46 cells, N = 17 mice; p = 1.35 × 10⁻⁹, unpaired two-tailed t-test). **(D)** Rin as a function of axon length for the population of recorded neurons (n = 66 cells, N = 18 mice). **(E)** Rin values of neurons with AIS and without AIS from the same data displayed in (D) (No AIS: 146.0 ± 42.52 MΩ [SD], n = 20 cells, N = 13 mice; AIS: 104.9 ± 30.70 MΩ [SD], n = 46 cells, N = 17 mice; p = 5.28 × 10⁻⁵, unpaired two-tailed t-test). **(F)** RMP as a function of axon length for the population of recorded neurons (n = 66 cells, N = 18 mice). **(G)** RMP values of neurons with AIS and without AIS from the same data displayed in (F) (No AIS: -50.13 ± 1.96 mV [SD], n = 20 cells, N = 13 mice; AIS: -51.40 ± 2.60 mV [SD], n = 46 cells, N = 17 mice; p = 0.0555, unpaired two-tailed t-test). **(H)** Firing gain in control and after the application of 10 μM 4-AP for the population of neurons without AIS, defined by the phase plot profile. (Control: 1.41 ± 0.65 spikes/nA [SD]; 4-AP: 2.25 ± 0.39 spikes/nA [SD]; n =17 cells, N = 11 mice; p = 2 × 10⁻⁶, paired two-tailed t-test). **(I)** Firing gain in control and after the application of 4-AP (10 μM) for the population of neurons with AIS, defined by the phase plot profile. (Control: 4.72 ± 5.36 spikes/nA [SD]; 4-AP: 18.01 ± 17.90 spikes/nA [SD]; n = 18 cells, N = 7 mice; p = 0.0017, paired two-tailed t-test). Data in H and I was used to generate plot in Fig. 2B. Horizontal bars in C, E, G, H and I represent population averages.

**Supplementary Figure 3.**
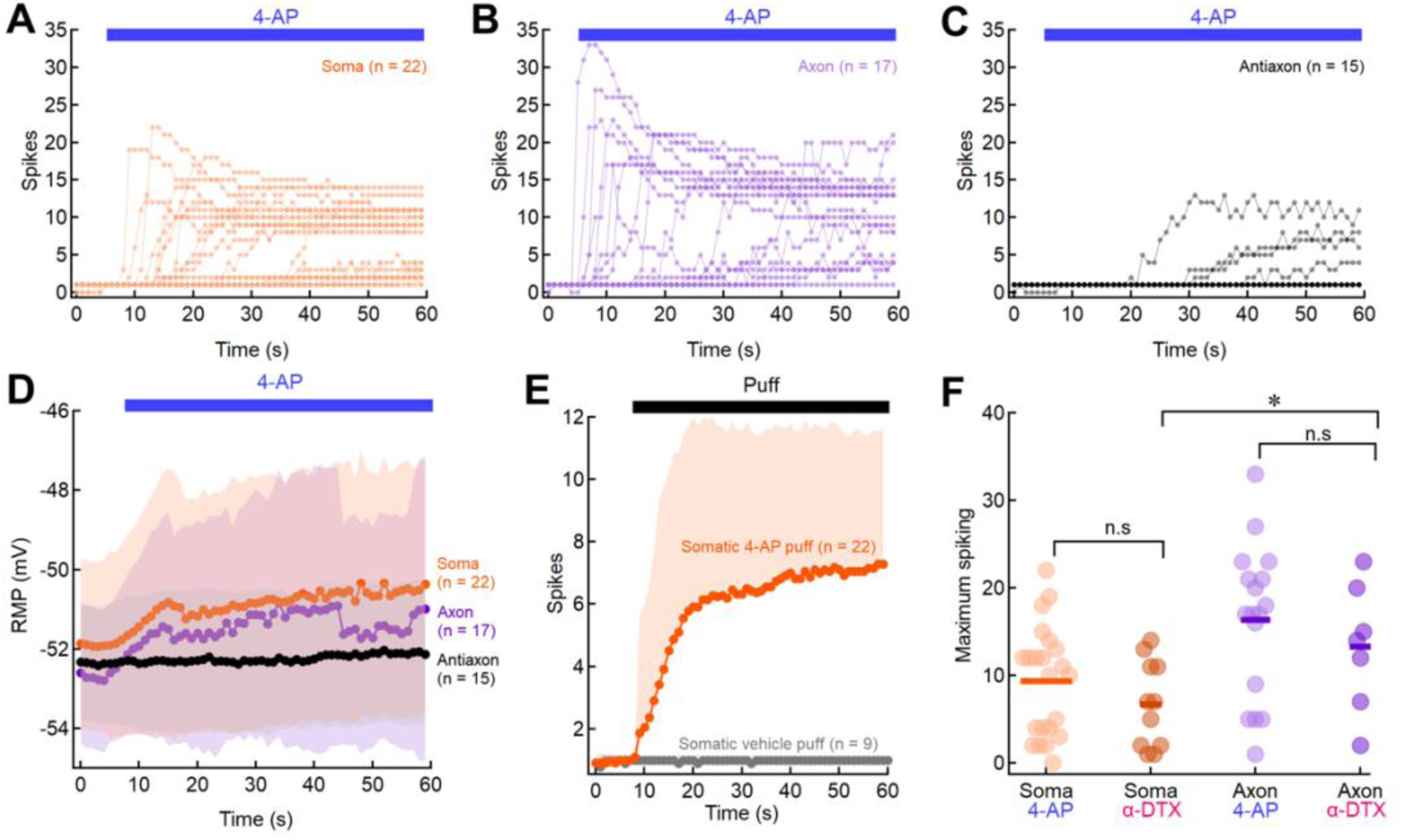
Additional information regarding Figure 2. Spike count per current pulse as a function of time for 4-AP puff applications at the soma **(A)**, axon **(B)**, and antiaxon **(C)** in the recorded neuronal population used to generate the plot shown in Figure 2E. Timing of 4-AP (100 μM) puff application is indicated by the horizontal bar at the top of each plot. **(D)** Plot of the RMP as a function of time for 4-AP puff applications at the soma (orange symbols, n = 22 cells, N = 9 mice), axon (purple symbols, n = 17 cells, N = 8 mice), and antiaxon (black symbols, n = 15 cells, N = 7 mice) in the recorded neuronal population. Timing of 4-AP (100 μM) application is indicated by the horizontal bar at the top of the plot. **(E)** Average plot of spike count per current pulse as a function of time for somatic puffing locations of 4-AP (orange symbols, n = 22 cells, N = 9 mice; same data as shown in Fig. 2E) and vehicle (aCSF; grey symbols, n = 9 cells, N = 3 mice). Timing of puff application is indicated by the horizontal bar at the top of the plot. **(F)** Plot of the maximum spike count per current pulse for each tested neuron during 4-AP (100 μM, same data as shown in Fig. 2 F) and α-DTX (2 μM) puffing applications (Somatic puff, 4-AP: 9.36 ± 6.25 spikes [SD], n = 22 cells, N = 9 mice; α-DTX: 6.73 ± 4.92 spikes [SD], n = 11 cells, N = 5 mice; p = 0.2320, unpaired two-tailed t-test. Axonal puff, 4-AP: 16.35 ± 8.72 spikes [SD], n = 17 cells, N = 8 mice; α-DTX: 13.29 ± 7.20 spikes [SD], n = 7 cells, N = 3 mice; p = 0.4212, unpaired, two-tailed t-test. α-DTX, soma vs. axon: p = 0.035, unpaired, two-tailed t-test). Shaded areas in D and E represent SD. Horizontal bars in F represent population averages.

**Supplementary Figure 4.**
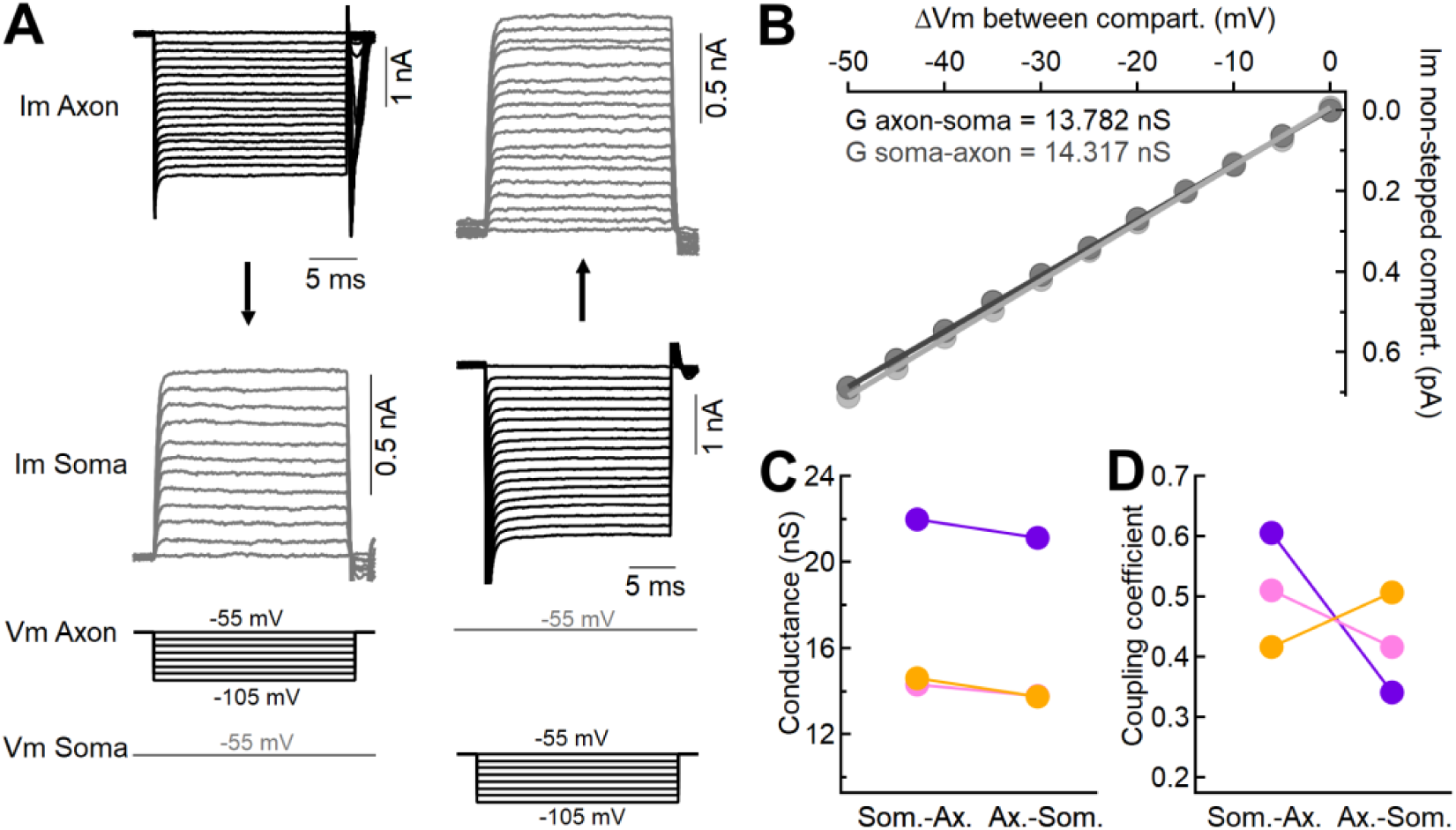
Electrotonic coupling between soma and the proximal axon. **(A)** Simultaneous voltage-clamp recordings from the soma and proximal axon of the same MesV neuron. Hyperpolarizing voltage steps were applied either to the axon (Vm axon, left) or to the soma (Vm soma, right), while the membrane potential of the other compartment was held constant. This protocol evoked inward membrane currents in the stepped compartment and corresponding outward currents in the non-stepped compartment (Im axon and Im soma). **(B)** Plot of the membrane current in the non-stepped compartment (Im non-stepped compart.) as a function of the voltage difference between compartments (ΔVm between compart.), obtained from the recordings shown in (A). Each dataset was fitted with a linear function, and the slope values, representing intercompartment conductance, are indicated (G axon-soma and G soma-axon). **(C)** Intercompartment conductance and **(D)** corresponding coupling coefficient values determined for each tested neuron in each direction (color-coded).

## REFERENCES

Abdollahi, N., Xie, Y.-F., Ratté, S., Prescott, S.A., 2025. K_V_ 1 Channels Enable Myelinated Axons to Transmit Spikes Reliably without Spiking Ectopically. J. Neurosci. 45, e1889242025. 10.1523/JNEUROSCI.1889-24.2025

Agmon-Snir, H., Carr, C.E., Rinzel, J., 1998. The role of dendrites in auditory coincidence detection. Nature 393, 268–272. 10.1038/30505

Alcami, P., 2018. Electrical Synapses Enhance and Accelerate Interneuron Recruitment in Response to Coincident and Sequential Excitation. Front. Cell. Neurosci. 12, 156. 10.3389/fncel.2018.00156

Alcami, P., Marty, A., 2013. Estimating functional connectivity in an electrically coupled interneuron network. Proceedings of the National Academy of Sciences of the United States of America 110, E4798–E4807. 10.1073/pnas.1310983110

Baker, R., Llinás, R., 1971. Electrotonic coupling between neurones in the rat mesencephalic nucleus. The Journal of Physiology 212, 45–63. 10.1113/jphysiol.1971.sp009309

Bean, B.P., 2007. The action potential in mammalian central neurons. Nature Reviews Neuroscience 8, 451–465. 10.1038/nrn2148

Bekkers, J.M., Delaney, A.J., 2001. Modulation of Excitability by α-Dendrotoxin-Sensitive Potassium Channels in Neocortical Pyramidal Neurons. J. Neurosci. 21, 6553–6560. 10.1523/JNEUROSCI.21-17-06553.2001

Bennett, M.V.L., 1966. Physiology of electrotonic junctions. Ann NY Acad Sci 137, 509–539. 10.1111/j.1749-6632.1966.tb50178.x

Bennett, M.V.L., Zukin, R.S., 2004. Electrical Coupling and Neuronal Synchronization in the Mammalian Brain. Neuron 41, 495–511. 10.1016/S0896-6273(04)00043-1

Carr, C., Konishi, M., 1990. A circuit for detection of interaural time differences in the brain stem of the barn owl. J. Neurosci. 10, 3227–3246. 10.1523/JNEUROSCI.10-10-03227.1990

Connors, B.W., Long, M.A., 2004. ELECTRICAL SYNAPSES IN THE MAMMALIAN BRAIN. Annual Review of Neuroscience 27, 393–418. 10.1146/annurev.neuro.26.041002.131128

Curti, S., Davoine, F., Dapino, A., 2022. Function and Plasticity of Electrical Synapses in the Mammalian Brain: Role of Non-Junctional Mechanisms. Biology 11, 81. 10.3390/biology11010081

Curti, S., Hoge, G., Nagy, J.I., Pereda, A.E., 2012. Synergy between electrical coupling and membrane properties promotes strong synchronization of neurons of the mesencephalic trigeminal nucleus. Journal of Neuroscience 32, 4341–4359. 10.1523/JNEUROSCI.6216-11.2012

Curti, S., O’Brien, J., 2016. Characteristics and plasticity of electrical synaptic transmission. BMC Cell Biology 17. 10.1186/s12860-016-0091-y

Dapino, A., Davoine, F., Curti, S., 2023. D-type K+ current rules the function of electrically coupled neurons in a species-specific fashion. Journal of General Physiology 155, e202313353. 10.1085/jgp.202313353

Davoine, F., Curti, S., 2019. Response to coincident inputs in electrically coupled primary afferents is heterogeneous and is enhanced by H-current (IH) modulation. Journal of Neurophysiology 122, 151–175. 10.1152/jn.00029.2019

Del Negro, C.A., Chandler, S.H., 1997. Physiological and theoretical analysis of K+ currents controlling discharge in neonatal rat mesencephalic trigeminal neurons. Journal of Neurophysiology 77, 537–553. 10.1152/jn.1997.77.2.537

Dessem, D., Taylor, A., 1989. Morphology of jaw-muscle spindle afferents in the rat. J of Comparative Neurology 282, 389–403. 10.1002/cne.902820306

Dodson, P.D., Barker, M.C., Forsythe, I.D., 2002. Two Heteromeric Kv1 Potassium Channels Differentially Regulate Action Potential Firing. J. Neurosci. 22, 6953–6961. 10.1523/JNEUROSCI.22-16-06953.2002

Florez-Paz, D., Bali, K.K., Kuner, R., Gomis, A., 2016. A critical role for Piezo2 channels in the mechanotransduction of mouse proprioceptive neurons. Sci Rep 6, 25923. 10.1038/srep25923

Fortin, S.M., Chen, J., Grill, H.J., Hayes, M.R., 2021. The Mesencephalic Trigeminal Nucleus Controls Food Intake and Body Weight via Hindbrain POMC Projections. Nutrients 13, 1642. 10.3390/nu13051642

Galarreta, M., Hestrin, S., 2001. Spike Transmission and Synchrony Detection in Networks of GABAergic Interneurons. Science 292, 2295–2299. 10.1126/science.1061395

Getting, P.A., 1973. Modification of Neuron Properties by Electrotonic Synapses. I. Input Resistance, Time Constant, and Integration. J. Physiol. 10.1152/jn.1974.37.5.846

Getting, P.A., Willows, A.O., 1974. Modification of neuron properties by electrotonic synapses. II. Burst formation by electrotonic synapses. Journal of Neurophysiology 37, 858–868. 10.1152/jn.1974.37.5.858

Gottlieb, S., Taylor, A., Bosley, M.A., 1984. The distribution of afferent neurones in the mesencephalic nucleus of the fifth nerve in the cat. J of Comparative Neurology 228, 273–283. 10.1002/cne.902280212

Haghdoust, H., Janahmadi, M., Behzadi, G., 2007. Physiological role of dendrotoxin-sensitive K+ channels in the rat cerebellar Purkinje neurons. Physiol Res 56, 807–813. 10.33549/physiolres.931041

Higgs, M.H., Spain, W.J., 2011. Kv1 channels control spike threshold dynamics and spike timing in cortical pyramidal neurones. J Physiol 589, 5125–5142. 10.1113/jphysiol.2011.216721

Inda, M.C., DeFelipe, J., Muñoz, A., 2006. Voltage-gated ion channels in the axon initial segment of human cortical pyramidal cells and their relationship with chandelier cells. Proc. Natl. Acad. Sci. U.S.A. 103, 2920–2925. 10.1073/pnas.0511197103

Joris, P.X., Smith, P.H., Yin, T.C.T., 1998. Coincidence Detection in the Auditory System. Neuron 21, 1235–1238. 10.1016/S0896-6273(00)80643-1

Joseph, A.W., Hyson, R.L., 1993. Coincidence detection by binaural neurons in the chick brain stem. Journal of Neurophysiology 69, 1197–1211. 10.1152/jn.1993.69.4.1197

Khakh, B.S., Henderson, G., 1998. Hyperpolarization-activated cationic currents (*I*_h_) in neurones of the trigeminal mesencephalic nucleus of the rat. The Journal of Physiology 510, 695–704. 10.1111/j.1469-7793.1998.00695.x

Kole, M.H.P., Stuart, G.J., 2012. Signal Processing in the Axon Initial Segment. Neuron 73, 235–247. 10.1016/j.neuron.2012.01.007

Kuba, H., 2003. Evaluation of the limiting acuity of coincidence detection in nucleus laminaris of the chicken. The Journal of Physiology 552, 611–620. 10.1113/jphysiol.2003.041574

Kuba, H., Koyano, K., Ohmori, H., 2002. Development of membrane conductance improves coincidence detection in the nucleus laminaris of the chicken. The Journal of Physiology 540, 529–542. 10.1113/jphysiol.2001.013365

Kuba, H., Yamada, R., Fukui, I., Ohmori, H., 2005. Tonotopic Specialization of Auditory Coincidence Detection in Nucleus Laminaris of the Chick. J. Neurosci. 25, 1924–1934. 10.1523/JNEUROSCI.4428-04.2005

Lazarov, N., Pilgrim, C., 1997. Localization of D1 and D2 dopamine receptors in the rat mesencephalic trigeminal nucleus by immunocytochemistry and in situ hybridization. Neuroscience Letters 236, 83–86. 10.1016/S0304-3940(97)00761-1

Lazarov, N.E., 2002. Comparative analysis of the chemical neuroanatomy of the mammalian trigeminal ganglion and mesencephalic trigeminal nucleus. Prog Neurobiol 66, 19–59. 10.1016/s0301-0082(01)00021-1

Lazarov, N.E., Chouchkov, C.N., 1995. Serotonin-containing projections to the mesencephalic trigeminal nucleus of the cat. The Anatomical Record 241, 136–142. 10.1002/ar.1092410118

Liem, R.S.B., Copray, J.C.V.M., Van Der Want, J.J.L., 1997. Dopamine-immunoreactivity in the rat mesencephalic trigeminal nucleus: an ultrastructural analysis. Brain Research 755, 319–325. 10.1016/S0006-8993(97)00124-8

Liem, R.S.B., Copray, J.C.V.M., Van Willigen, J.D., 1991. Ultrastructure of the Rat Mesencephalic Trigeminal Nucleus. Cells Tissues Organs 140, 112–119. 10.1159/000147045

Lorincz, A., Nusser, Z., 2008. Cell-Type-Dependent Molecular Composition of the Axon Initial Segment. J. Neurosci. 28, 14329–14340. 10.1523/JNEUROSCI.4833-08.2008

Luo, P.F., Wang, B.R., Peng, Z.Z., Li, J.S., 1991. Morphological characteristics and terminating patterns of masseteric neurons of the mesencephalic trigeminal nucleus in the rat: An intracellular horseradish peroxidase labeling study. J. Comp. Neurol. 303, 286–299. 10.1002/cne.903030210

Mathews, P.J., Jercog, P.E., Rinzel, J., Scott, L.L., Golding, N.L., 2010. Control of submillisecond synaptic timing in binaural coincidence detectors by Kv1 channels. Nat Neurosci 13, 601–609. 10.1038/nn.2530

Morquette, P., Lavoie, R., Fhima, M.-D., Lamoureux, X., Verdier, D., Kolta, A., 2012. Generation of the masticatory central pattern and its modulation by sensory feedback. Progress in Neurobiology 96, 340–355. 10.1016/j.pneurobio.2012.01.011

Overholt, E., Rubel, E., Hyson, R., 1992. A circuit for coding interaural time differences in the chick brainstem. J. Neurosci. 12, 1698–1708. 10.1523/JNEUROSCI.12-05-01698.1992

Perez Velazquez, J.L., Carlen, P.L., 2000. Gap junctions, synchrony and seizures. Trends in Neurosciences 23, 68–74. 10.1016/S0166-2236(99)01497-6

Rela, L., Szczupak, L., 2003. Coactivation of Motoneurons Regulated by a Network Combining Electrical and Chemical Synapses. J. Neurosci. 23, 682–692. 10.1523/JNEUROSCI.23-02-00682.2003

Scott, L.L., Mathews, P.J., Golding, N.L., 2005. Posthearing Developmental Refinement of Temporal Processing in Principal Neurons of the Medial Superior Olive. J. Neurosci. 25, 7887–7895. 10.1523/JNEUROSCI.1016-05.2005

Shu, Y., Duque, A., Yu, G., Haider, B., McCormick, D.A., 2007a. Properties of action-potential initiation in neocortical pyramidal cells: Evidence from whole cell axon recordings. Journal of Neurophysiology 97, 746–760. 10.1152/jn.00922.2006

Shu, Y., Yu, Y., Yang, J., McCormick, D.A., 2007b. Selective control of cortical axonal spikes by a slowly inactivating K + current. Proc. Natl. Acad. Sci. U.S.A. 104, 11453–11458. 10.1073/pnas.0702041104

Spitzer, M.W., Semple, M.N., 1995. Neurons sensitive to interaural phase disparity in gerbil superior olive: diverse monaural and temporal response properties. Journal of Neurophysiology 73, 1668–1690. 10.1152/jn.1995.73.4.1668

Takahashi, T., Shirasu, Masayoshi, Shirasu, Mari, Kubo, K.Y., Onozuka, M., Sato, S., Itoh, K., Nakamura, H., 2010. The locus coeruleus projects to the mesencephalic trigeminal nucleus in rats. Neuroscience Research 68, 103–106. 10.1016/j.neures.2010.06.012

Tanaka, S., Wu, N., Hsaio, C.F., Turman, J., Chandler, S.H., 2003. Development of inward rectification and control of membrane excitability in mesencephalic V neurons. Journal of Neurophysiology 89, 1288–1298. 10.1152/jn.00850.2002

Trigo, F.F., Alcamí, P., Curti, S., 2025. Functional interaction of electrical coupling and H-current and its putative impact on inhibitory transmission. Neuroscience 574, 13–20. 10.1016/j.neuroscience.2025.03.051

Verdier, D., Lund, J.F., Kolta, A., 2004. Synaptic inputs to trigeminal primary afferent neurons cause firing and modulate intrinsic oscillatory activity. Journal of Neurophysiology 92, 2444–2455. 10.1152/jn.00279.2004

Veruki, M.L., Hartveit, E., 2002. AII (Rod) Amacrine Cells Form a Network of Electrically Coupled Interneurons in the Mammalian Retina. Neuron 33, 935–946. 10.1016/S0896-6273(02)00609-8

Weinberg, E., 1928. The mesencephalic root of the fifth nerve. A comparative anatomical study. J of Comparative Neurology 46, 249–405. 10.1002/cne.900460202

Yin, T.C., Chan, J.C., 1990. Interaural time sensitivity in medial superior olive of cat. Journal of Neurophysiology 64, 465–488. 10.1152/jn.1990.64.2.465

